# Distant Residues Modulate Conformational Opening in SARS-CoV-2 Spike Protein

**DOI:** 10.1101/2020.12.07.415596

**Authors:** Dhiman Ray, Ly Le, Ioan Andricioaei

## Abstract

Infection by SARS-CoV-2 involves the attachment of the receptor binding domain (RBD) of its spike proteins to the ACE2 receptors on the peripheral membrane of host cells. Binding is initiated by a down-to-up conformational change in the spike protein, the change that presents the RBD to the receptor. To date, computational and experimental studies that search for therapeutics have concentrated, for good reason, on the RBD. However, the RBD region is highly prone to mutations, and is therefore a hotspot for drug resistance. In contrast, we here focus on the correlations between the RBD and residues distant to it in the spike protein. This allows for a deeper understanding of the underlying molecular recognition events and prediction of the highest-effect key mutations in distant, allosteric sites, with implications for therapeutics. Also, these sites can appear in emerging mutants with possibly higher transmissibility and virulence, and pre-identifying them can give clues for designing pancoronavirus vaccines against future outbreaks. Our model, based on time-lagged independent component analysis (tICA) and protein graph connectivity network, is able to identify multiple residues that exhibit long-distance coupling with the RBD opening. Residues involved in the most ubiquitous D614G mutation and the A570D mutation of the highly contagious UK SARS-CoV-2 variant are predicted *ab-initio* from our model. Conversely, broad spectrum therapeutics like drugs and monoclonal antibodies can target these key distant-but-conserved regions of the spike protein.

**Significance Statement:** The novel coronavirus (SARS-CoV-2) pandemic resulted in the largest public health crisis in recent times. Significant drug design effort against SARS-CoV-2 is focused on the receptor binding domain (RBD) of the spike protein, although this region is highly prone to mutations causing therapeutic resistance. We applied deep data analysis methods on all-atom molecular dynamics simulations to identify key non-RBD residues that play a crucial role in spike-receptor binding and infection. textcol-orredBecause the non-RBD residues are typically conserved across multiple coronaviruses, they can be targeted by broad spectrum antibodies and drugs to treat infections from new strains that might appear during future epidemics.

---

**T**he COVID-19 pandemic continues its spread, with more than 160 million confirmed cases worldwide, including 4.1 million deaths by the end of July 2021, according to the World Health Organization. The etiological agent, SARS-CoV-2, is a member of the *Coronaviridae* family, which includes SARS-CoV-1 (2002-2004) and MERS-CoV (since 2012), viruses with which SARS-CoV-2 has a sequence identity of 79.6% and 50%, respectively (1, 2). Expressed on the surface of SARS-CoV-2, the spike (S) protein plays a crucial role in infection. It binds to the host angiotensin-converting enzyme 2 (ACE2) through the S protein’s receptor-binding domain (RBD), thereby facilitating viral entry into host cells (3, 4). Therefore, the spike protein is a preponderant target for inhibitors that impede SARS-CoV-2 infection. Assessing the genomic variability of SARS-CoV-2 reveals a moderate mutation rate compared to other RNA viruses, around 1.12 × 10^-3^ nucleotide substitutions/site/year (5), (but much larger than DNA viruses (6)); the SARS-CoV-2 mutation rate is at the same level as SARS-CoV-1 (7). Significantly, the spike protein has been demonstrated to be particularly susceptible to acquiring mutations (5, 8–10). More specifically, a study analyzing 10,022 SARS-CoV-2 genomes from 68 countries revealed 2969 different missense variants, with 427 variants in the S protein alone (5). This suggests a strong propensity to form new strains with higher virulence and more complicated epidemiology; the dominant D614G mutation and the recent B.1.1.7 mutant are notable examples (11). Spike protein variability can thus render currently existing therapeutic agents ineffective in combating SARS-CoV-2 and other probable SARS epidemics in the future. It therefore is of fundamental value to understand, in microscopic detail, the role of spike mutations to the structural dynamics that triggers infection.

Large scale screening of therapeutic molecules and antibodies are underway, aiming to target the spike protein and consequently to prevent infection. Most of the experimental (12–15) and computational (16–18) efforts for inhibitor design focus on the receptor binding domain (RBD), despite the fact that this region is highly mutation-prone (see, for example, Verkhivker (19), Spinello et al. (20) and our own sequence alignment study in the SI Appendix) and can cause resistance to therapeutics. For instance, the mutations in the RBD observed in the emerging viral lineage in South Africa resulted in up to a ten-fold reduction in the neutralization capacity of conventional antibody therapy (21).

However, domains in the spike other than the RBD are also possible targets for inhibition. The human immune system started generating antibodies specific to residues outside the RBD even at the early stages of the pandemic. Liu et al. extracted, from infected patients, multiple COVID-19 neutralizing antibodies, a fraction of which bind to non-RBD epitopes of the S-protein, such as the N-terminal domain (NTD) (22). Moreover, a separate group of antibodies, present in the blood-stream of uninfected humans (particularly, and importantly, in children), were observed to bind specifically to the S2-stem region. This structure has virtually identical sequence in all coronavirus strains, including the ones causing common cold (23). It is the presence of such universal antibodies, selective to the conserved regions of the spike, that is hypothesized to cause the absence of severe infections in children. These observations motivate an effort towards designing small-molecule drugs and antibodies targeted towards residues far from the RBD in the 3D structure which, consequently, are less prone to mutation.

A number of studies explored druggable hotspots in the spike protein that modulate, via allostery, ACE2 binding (20, 24–26). A concurrent computational study established that RBD mutations appearing in the new strains of SARS-CoV-2 can, via allosteric effects, increase binding affinity ^of^ the spike to the human ACE2 receptor as well as impair the binding interactions of neutralizing antibodies (20). Hydrogen deuterium exchange mass spectrometry (HDXMS) experiments revealed the dampening and the increase of conformational motion at the stalk region and the proteolytic cleavage site respectively, upon RBD binding to the ACE2 receptor (26). Although this is a noteworthy approach towards targeting non-RBD residues, the S-ACE2 bound complex, once formed, already can initiate infection. So inhibitors which bind to the ACE2-bound spike are only partially effective in preventing viral entry. We, therefore, focus our study on the effect of distant residues on the dynamics of the structural transition in the spike protein leading to the RBD-up conformation, i.e., *before* it becomes posed for binding (see Fig 1). This is because RBD opening plays an obligatory role in the infection by displaying the RBD to the ACE2 receptor. (27–31). Therapeutics inhibiting this structural transition can therefore prevent ACE2 binding altogether, providing a higher degree of barrier towards the infection.

**Fig. 1.**
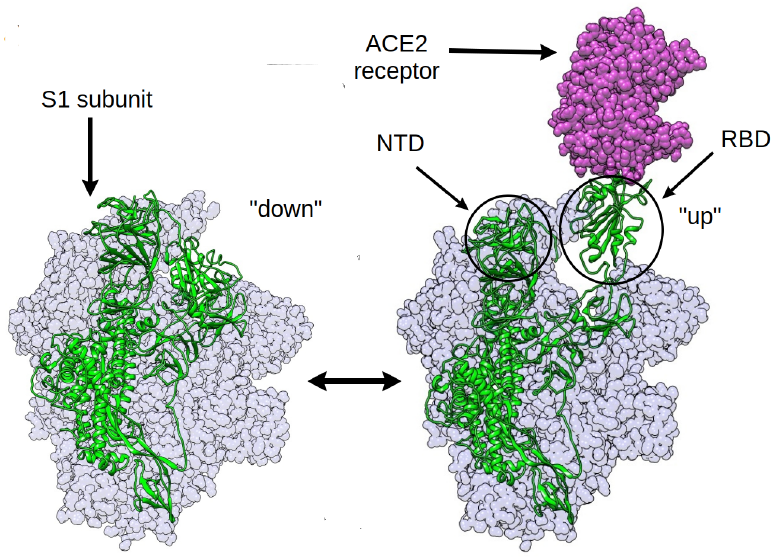
The RBD opening transition in the SARS-CoV-2 spike protein: the RBD of the chain shown in green is undergoing the *down* to *up* transition leading to the binding of the human ACE2 receptor.

The down-to-up transition of the S-protein RBD has been subjected to extensive cryogenic electron microscopy (cryoEM) studies that elucidated the atomic resolution structures of the closed (PDB ID: 6VXX (32), 6XR8 (33)), partially open (PDB ID: 6VSB (34)) and fully open (PDB ID: 6VYB (32)) states. Attempts have been made to study the dynamics of RBD opening using force-field based classical molecular dynamics (MD) simulation (35, 36). However, the direct observation of an RBD opening transition is beyond the scope of atomistic MD simulation, primarily because of the large size of the system and the long timescales involved. Yet, combining multiple short trajectories totaling up to 1.03 ms, the Folding@Home project could resolve various intermediates of the RBD opening transition (36). Alternatively, Gur et al. used steered molecular dynamics in combination with unbiased MD simulations to uncover the mechanism of the conformational opening process (30). They obtained a free energy landscape and delineated the effect of salt bridges on the transition. Moreover, structure-based coarse-grained modeling could be employed to explore the conformational landscape of multiple configurations of the spike trimer (1-up-2-down, 2-up-1-down, 3-up and 3-down) (37), and, in a separate study, to identify allosteric communication pathways in the spike protein (19). Despite providing qualitative insight, the absence of explicit solvent and atomistic detail and the exclusion of the glycan-shield (which plays dynamic roles beyond shielding itself (35, 38)) obscured a quantitative mechanistic picture of the conformational opening transition.

We took a three-pronged approach to identify the distant residues that show correlated motion coupled to RBD opening and closing. First, as an initial exploration, we pulled the RBD of the closed state at non-equilibrium, to generate a putative open structure. Using potential of mean force (PMF) calculations, obtaining an non-equilibrium work profile, this structure, as expected, lead to a value higher than the equilibrium free energy (20 kcal/mol, see Fig S1 in SI Appendix). This is akin to a single-molecule pulling experiment (39). However, we emphasize that we do not use the reaction coordinate (RC) to derive any quantitative data. Given the artificiality of the result obtained by following a single degree of freedom, we resorted to the use of the more sophisticated tICA coordinates to capture the motion. We then designed and employed a novel way to identify residues important for the conformational change by quantifying the correlation of the backbone torsion angles of the protein with the slowest degrees of freedom, representing the down-to-up transition of the RBD. Thirdly, we took an alternative route to study allosteric connections by constructing a protein graph connectivity network model that uses the mutual information metric computed from MD trajectories. Taken together, the three approaches resulted in the prediction of a handful of residues in non-RBD regions of the S protein that play a crucial role in the conformational rearrangements of the spike. These residues suggest possible future mutational hotspots, as well as targets for designing inhibitors that can reduce the flexibility of the RBD, leading to reduced receptor binding capability. In fact, the most ubiquitous spike protein mutation to date, the D614G (40), and the A570D mutation in the recently emerged highly contagious UK SARS-CoV-2 strain (41), both appeared among our predicted set of residues, the latter having been discovered after the completion of our calculations.

## Results

### Correlation between RBD-opening and backbone dihedral angle

Multiple unbiased trajectories were propagated from different regions of the S-protein RBD-opening conformational space, priorly explored by steered MD and umbrella sampling (details in Supporting Information (SI) Appendix). Three of those trajectories were assigned as the closed, partially open, and fully open state, based on the position of free energy minima along the umbrella sampling reaction coordinate (cf. Fig. S1 in SI Appendix). The stability of these three conformations were ensured by inspection of the trajectories. The cumulative simulation data was projected onto a feature space composed of pairwise distances between residues from RBD and from other parts of the spike near the RBD (details in SI Appendix); these distances increase during the down-to-up transition. The projected trajectory data were subjected to principal component analysis (PCA) (42) and time-lagged independent component analysis (tICA) (43–45). The former method calculates the degrees of freedom with the highest variance in the system, while the latter obtains the ones with the longest timescale. These methods quantify large conformation changes in complex bio-molecules. The goal of performing PCA and tICA is to find out one or two coordinates which best describe the RBD opening motion. As one continuous trajectory hardly samples the transition event, multiple short trajectories spanning a large range of the configuration space were used (46).

The first principal component (PC) and the first time-lagged independent component (tIC), obtained from PCA and, respectively, from tICA analysis could both distinguish the closed and open states, although the partially open and fully open states could not be distinguished within the first two PCs (Fig. S2 in SI Appendix). Yet, the projections of the open- and closed-state trajectories along the first two principal components are in agreement with the results of previous long multi-microsecond spike protein simulations (35). Because of the clear distinction between the closed, open and the intermediate, partially open states (Fig. 2b), we chose the first two time-lagged independent components (tICs) for our subsequent analysis.

**Fig. 2.**
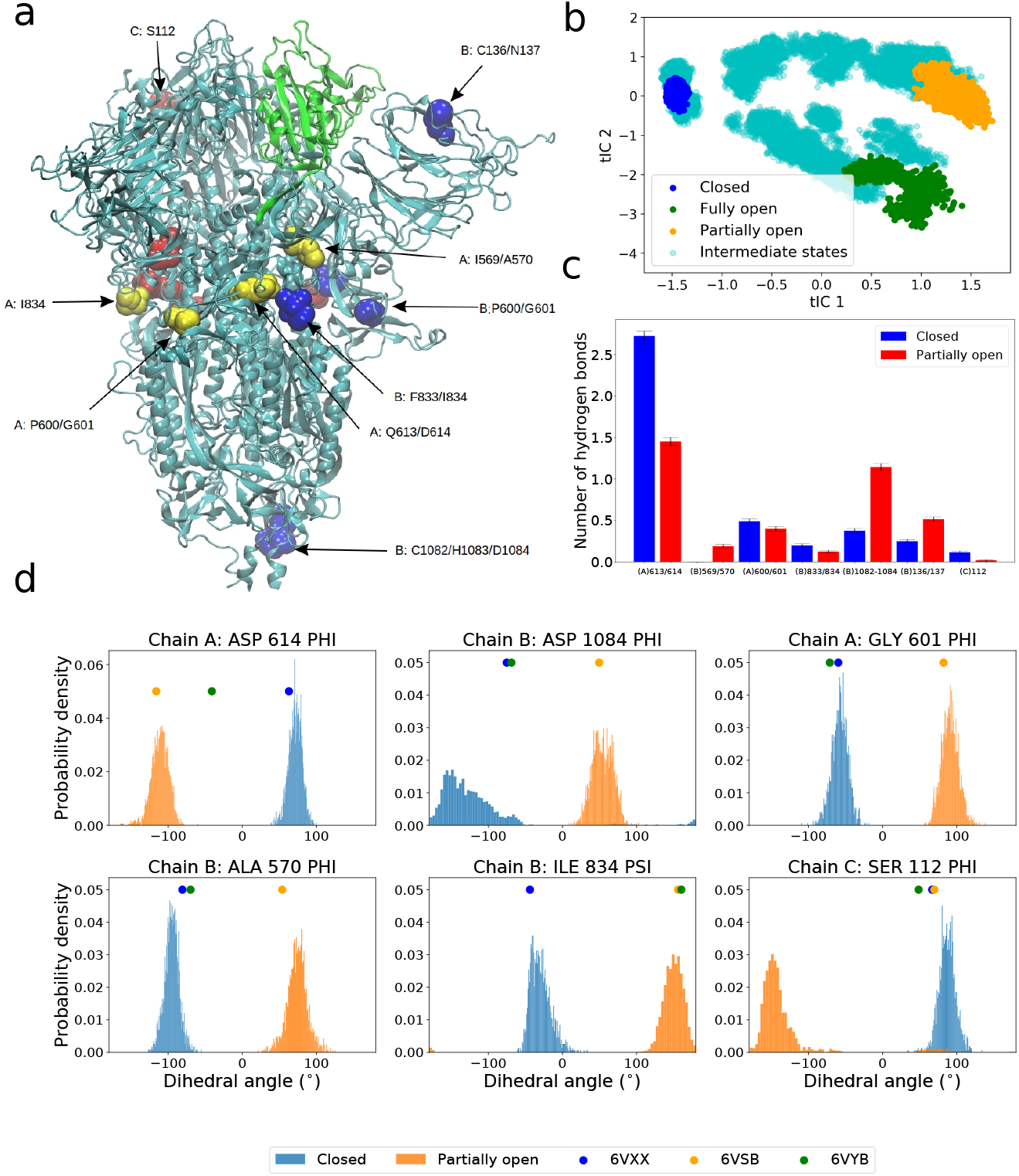
(a) Structure of spike protein with the residues in RBD shown in green color. Non-RBD residues strongly correlated with the RBD opening motion are represented by spheres (color code: chain A: yellow, chain B: blue, chain C: red). The RBD of chain A is performing *down-to-up* conformational change. (b) The projection of all unbiased trajectories along the two slowest degrees of freedom (tICs) obtained from tICA analysis. (c) Average number of hydrogen bonds for the highest correlated residues/residue-pairs (Table 1) in the closed and the partially open state. (d) Normalized distribution of representative backbone dihedral angles strongly correlated with tIC 1 and tIC 2 (Table 1). The distributions are calculated from closed and partially open state trajectories. The corresponding values of the dihedral angles in the PDB structures are marked in the plot for reference.

Large scale conformational changes in proteins are, at a fundamental level, stemming from complex combinations of transitions between various states of the protein-backbone torsional angles *ϕ* and *ψ*. These combinations add up to global displacements, which set the timescale for internal friction (47) and gauge the paradoxically large number of conformational states accessible to a protein as it folds (48). On one hand, concerted transitions of the backbone torsions typically lead to large scale motion. On the other hand, exponential divergence in nonlinear dynamical systems (49) is such that only certain dihedrals are likely to predominantly effect the conformational changes. We therefore hypothesize that there exist specific residues in the spike protein, for which the transition in backbone dihedral states result in the opening of the RBD. Moreover, we conjecture that, given the significant mutation rate, and because of the selection pressure on the virus, the residues with large impact in collective motions that facilitate infectivity are likely to be selected in spike mutants. To test this hypothesis we calculated the Pearson correlation coefficients of the sines and cosines of all the *ϕ* and *ψ* backbone torsion angles of all the residues with the first two tICs. The magnitude of the correlation is found to be significantly large only for a handful of torsion angles, whereas the majority show near-zero correlation (Fig. S3 in SI Appendix).

We defined a metric called “correlation score” (*CS*) for each torsion angle in each residue. The value of the metric is computed as:

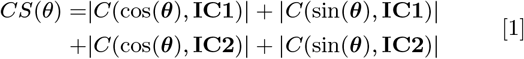

where ***θ***, **IC1** and **IC2** are the vectors containing the time series of the angle *θ*, the tIC1 and tIC2 respectively, and *C*(***x*, *y***) is the Pearson correlation coefficient of datasets **x** and **y**. The *CS* metric can take values from 0 to 4 and a higher value indicates that the particular torsion angle shows a highly correlated (or anticorretated) motion with the slowest conformational change which in this case the RBD transition. We avoided summing over the Φ and Ψ angles of the same residue, or over residues of different protein chains as it might average out the contributions from each angle and consequently obscure the process of specifying the role of each individual residue in the conformational transition.

We sorted the residues based on the *CS* scores of their torsion angles and a list is provided in the SI. The highest values of correlation scores are shown primarily by pairs of consecutive residues, with *ψ* of the first and the *ϕ* of the second residue, as depicted in Table 1. This suggests that two consecutive torsion angles in certain regions of the protein are highest correlated with the RBD opening motion. Most dihedrals belonged to residues in the loop structure joining the RBD with the S2 stem, as this region is a hinge for the opening of the RBD. This correlation does not exclude causation, since change in the conformational state for two subsequent torsion angles can induce crankshaft motion (50) in the backbone which, propagating along the chains, leads to a change in protein structure.

**Table 1.** A list of residues for which the backbone torsional angles are strongly correlated with the first two tIC components. The correlation score for each of the dihedral angles (see text and SI Appendix) is also reported.

| Residue | Chain | Predominant angle | Correlation score (*CS*) |
| --- | --- | --- | --- |
| Gln613 | A, B, C | $\psi$ | 2.62, 2.61, 2.59 |
| Asp614 | A, B, C | $\phi$ | 2.56, 2.54, 2.46 |
| Cys1082 | B | $\phi$ | 2.25 |
| His1083 | B | $\psi$ | 2.54 |
| Asp1084 | B | $\phi$ | 2.49 |
| Pro600 | A, B, C | $\psi$ | 1.56, 1.48, 1.52 |
| Gly601 | A, B, C | $\phi$ | 2.37, 2.31, 2.33 |
| Ile569 | A, B, C | $\psi$ | 2.40, 2.49, 2.46 |
| Ala570 | A, B, C | $\phi$ | 1.56, 2.22, 2.16 |
| Cys136 | B | $\psi$ | 1.42 |
| Asn137 | B | $\psi, \phi$ | 1.70, 1.80 |
| Phe833 | B | $\psi$ | 2.37 |
| Ile834 | A, B, C | $\psi$ | 1.73, 2.40, 2.12 |
| | A | $\phi$ | 1.19 |
| Ser112 | C | $\psi, \phi$ | 1.67, 2.23 |

The distribution of some of the dihedral angles, with highest *CS* scores, in the closed and partially open states, are depicted in Fig. 2d. Similar plots for all torsion angles with *CS* > 2.0 are provided in the SI Appendix. We compared the dihedral angle distribution only beween the closed and the open states as the significance of the artificially prepared fully open structure should not be overemphasized. This structure, generated from the closed state using steered molecular dynamics and umbrella sampling (see SI Appendix), is only an approximation for a more exact RBD up structure that binds the ACE2 receptor. In literature the structure with PBD ID: 6VSB is sometimes referred to as the open conformation (35), which, in this work, we refer to as the “partially open” state. So, in the rest of the paper, when we attempt to compare the behaviour of a chosen set of residues between closed and open states at an atomistic detail, we only include the closed and partially open configurations.

The fact, that the highly correlated residues follow a distinctly different distribution in the backbone torsion angle space (Fig 2d), indicates that a handful of non-RBD residues can play a pertinent role in the conformational change of the spike and, consequently, in the viral infection. Interestingly, the correlated torsion angles span over all three chains of the spike trimer, namely A (the one undergoing the RBD transition), B and C (Fig. 2a and Table 1), hinting at the potential role of inter-residue couplings ranging over long distances in presenting the RBD to the ACE2 receptor. The residues exhibiting highest correlation scores (Table 1), particularly Gln613, Asp614, Pro600, Gly601, Ile569 and Ala570, are present in the linker region joining the RDB with the S2 domain, which, as mentioned above, is the hinge for the opening motion that presents the RBD to the receptor (51). The Phe833 and Ile834 residues, although technically part of S2 domain, can significant impact the dynamics of the hinge or linker due to their proximity in 3D structure. Similar arguments are applicable for the NTD domain residues such as Cys136 and Asn137 from chain B and Ser112 from chain C, which are able to impact the RBD due to their structural proximity. Interestingly, residues near the stem region, including Cys1082, His1083, and Asp1084 appear in our list as strongly correlated and can potentially be used as a target for broad spectrum antibody or vaccine design targeting the stem region (52, 53).

When the virus mutates these particular residues in a way that increases its virulence, this increase stems from the propensity of the RBD to “flip” open and thereby increase ACE2 binding. As evidence, we highlight the example of the D614G mutation, which is already observed in numerous strains of the SARS-CoV-2 all over the world (10, 40). Cryo-EM studies have indicated that the D614G mutation is, by itself, capable of altering the the conformational dynamics of spike protein by stabilizing a RBD up state over the down conformation (54, 55). D614 is one of the top ranked residues predicted from our model for the potential to play a crucial role in RBD opening. A glycine residue has the least backbone-torsion barrier for conformational transition in *ϕ*-*ψ* space due to the absence of a side chain. Replacing an Asp residue, which has higher barriers to such transitions, with a glycine can increase the flexibility of the backbone, significantly impacting the probability of observing an RBD-up conformation. To understand if this can be the reason why this particular mutation was selected to become so widespread, a comparison of the geometric and “chemical” effects of Gly should be assessed. To this end, we performed additional simulations of the open and closed states of the D614G mutant. Indeed, our simulations of the D614G mutant spike indicate that, unlike the wild type system for which a significantly different dihedral angle distribution exists, there is no difference between the closed and the partially open configuration in terms of the torsion angle space explored by residue G614 (see Fig. S14 in SI Appendix). The glycine residue at 614 position also experienced different degrees of hydrogen bonding and electrostatic interactions (see below).

A wide range of spike protein mutant sequences have been characterized, each with varying degrees of abundance. A relatively rare mutation, A570V, resulted in a decrease of the overall stability of the spike protein in all three states, based on the FoldX empirical force field (10, 56). Free energy values (10) were obtained from only structural data and no dynamical information was considered in that study. Yet it is worth noting that the change in total and solvation free energies, due to this mutation, were substantially different for the closed and open states, resulting in a change in *△G* for RBD opening. But, as the side chains of Ala and Val are similar in terms of steric bulk, this mutation is unlikely to significantly impact RBD dynamics. As it likely did not increase the evolutionary advantage of the virus by increasing infectivity, this mutation, only occurring in one strain (10) so far, did not become as prevalent as D614G.

On the contrary, a A570D mutation is observed in the same residue in the newly emerged and highly infectious B.1.1.7 strain in the UK (41, 57, 58). This mutation is likely to play a pertinent role in infection as it replaces a hydrophobic amino acid with a charged one. This leads to a significant difference in the conformational dynamics of the 570 residue and consequently impacts the large scale RBD opening motion. Structural biology experiments have established that a mutation in the A570 residue alters the propensity of RBD opening by modulating the hydrophobic interaction of the hinge region with the S2 core (59). Coarse grained modeling studies explained this observation by noting that A570 is part of a regulatory switch that triggers the conformation change necessary for receptor binding (60).

The B.1.1.7 also shows a P681H mutation close to the highly correlated N679 residue predicted from our model (see Fig. S5 in SI Appendix). As this mutation replaces a structurally rigid proline residue, it can possibly impact the conformational space accessible to nearby residues, including N679.

Giving pause for thought, these results indicate that mutations in the highest correlated residues (Table 1) can in fact have significant physiological impact in changing the course of the pandemic. Therefore, we provide a list of residues (Table 1 and Fig. S3-S10 in SI Appendix), future mutations of which could impact RBD dynamics and consequently change the transmissibility or virulence of SARS-CoV-2. (See data availability statement for access to the raw correlation coefficient data for all residues.) Yet, care should be taken with assigning the predominant role in infection to a single, non-RBD domain residue in the UK variant; several other mutations are present that could modulate the binding affinity to the ACE2 receptor (particularly N501Y in the RBD-ACE2 binding interface). However, mutations outside the RBD can indeed play a key role in infection by disproportionately favoring an RBD “up” structure (52, 53, 59).

For a more detailed understanding we compared the average number of hydrogen bonds per residue group from Table 1 for the closed and the partially open trajectory (Fig 2c). We observed significant changes in the number of hydrogen bonds in residue groups: Q613/D614, I569/A570, C1082/H1083/D1084, and C136/N137 (Fig. 3). In the closed state, the carboxylate side-chain of D614 residue forms hydrogen bonds with K854 and T859 which are lost in the RBD up configuration. These hydrogen bonds will be absent in D614G mutant and likely reduce the energy cost of the conformational transition. Particularly the loss of hydrogen bond with T859 has been attributed to the higher stability of the RBD up structure in D614G mutant by Mansbach et al. (61). Our simulations also indicate that there is a loss of one hydrogen bond in the Q613/D614 residues going from RBD down to RBD up conformation in the WT spike. But such loss of hydrogen bonding is not observed in case of D614G mutant (see SI Appendix). On the contrary, formation of new hydrogen bonds are observed in the other three residue groups (Fig. 3) which can be enhanced or reduced by mutating the residues involved.

**Fig. 3.**
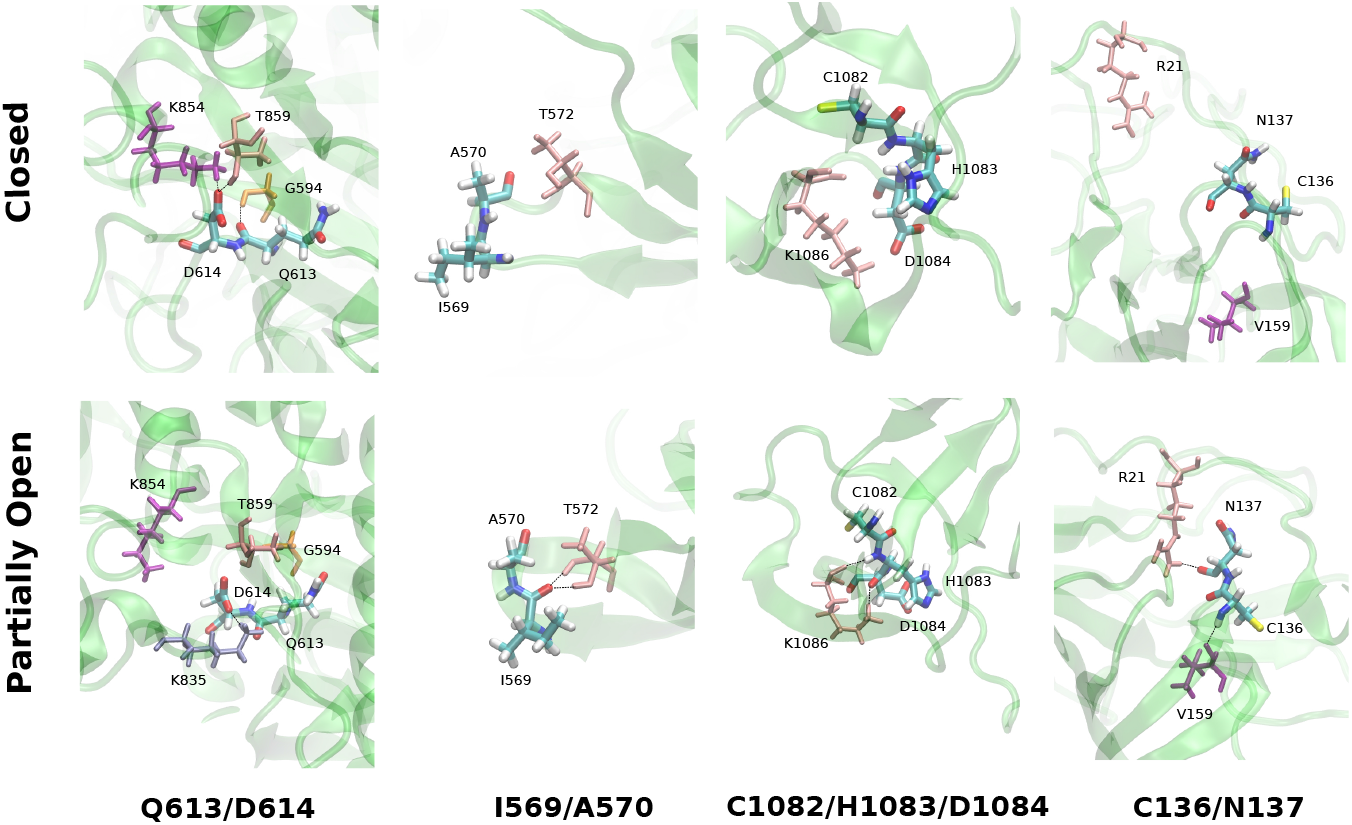
Representative snapshots of the hydrogen bonding pattern of some of the groups of residues from Table 1 and Fig. 1c. The upper panel corresponds to the closed state state and the lower panel shows the partially open state.

Additionally, the non-bonded interaction energies (electrostatic and van der Waals (vdW)) of the residues, from Table 1, differ significantly in the two conformations. Unsurprisingly, the D614 is energetically stabilized in the closed conformation in comparison to the open state due to additional hydrogen bonds. In the D614G mutant, from our analysis, this stabilization is significantly lower in comparison to the WT (see Fig. S15 and discussion in SI Appendix.) But residues P600/G601 are more stabilized in the open state in comparison to the closed state via favourable Coulomb and vdW interactions. Similar effect is observed in C136 for electrostatic energy but is somewhat compensated for by the opposite trend in vdW energy. A570 and N137 have lower electrostatic energy in the closed state despite having fewer hydrogen bonds. In short, non-RDB residues that experience different amount of non-bonded interaction with the rest of the protein or show different hydrogen bonding patterns in the “RBD down” and “RBD up” state, can impact the relative stability of those two conformations when mutated into residues with different properties.

### Mutual information and network model

A different approach to characterize the coupling between distant regions in a protein is to calculate the cross correlations between the positions of different residues in 3D space. This method is often used to study allosteric effects upon ligand binding (62–67). Conventional implementations compute the dynamic cross correlation map (DCCM) of the position vectors of the *C_α_* atoms (68). However, DCCM ignores correlated motions in orthogonal directions (67). This problem can be avoided by using a linear mutual information (LMI) based cross-correlation metric, which we use in the current study (62, 69). The cross-correlation matrix elements, *C_ij_*, are given by

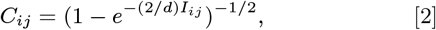

where *I_ij_* is the linear mutual information computed as

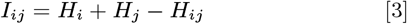

with *H* the Shannon entropy function:

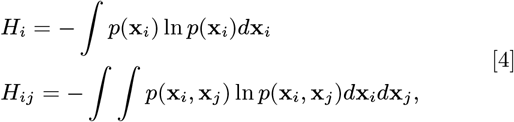

where, for two residues *i* and *j*, *X_i_*, *X_j_* are the 3-dimensional Cartesian vectors of atomic coordinates of the corresponding *C_α_* atom, whereas *p*(*x_i_*) and *p*(*x_i_*, *X_j_*) indicate, respectively, the marginal probability density for *X_i_* and the joint probability density of *x_i_* and *x_j_* (62, 69).

The change in cross-correlation between the apo and holo states of a protein is a gauge that traces allosteric communication in the protein by monitoring the changes in the local correlations between protein residues(62). The spike protein conformational change is not an allosteric process by strict definition as it does not involve the binding of an effector. But comparing the LMI cross correlations between the RBD-down and up states can help identify residues which behave differently in different protein conformations. More importantly the difference of correlation between the closed state and a small perturbed structure towards the conformational opening, can hint at the residues which gain or loose contact in the beginning stage of the opening transition and consequently initiating the large scale motion. So we assigned one of the unbiased trajectories as “slightly open”, indicative of an early stage structure in the pathway of opening. We included the unbiased trajectory, corresponding to this structure, in our subsequent analysis, in which we compared the correlation heat map of all residues in the closed form with the three open states, namely the partially open, fully open, and slightly open state, in order to understand the coupling between RBD opening and protein residue fluctuations.

Overall change in LMI correlation is clearly larger for the fully open state in comparison to the partially open state as evident from the higher appearance of reddish color (Fig. 4 a-d). Unsurprisingly, the change is largest for the RBD and proximal residues encompassing the N-terminal domain region (residue 100-300) in all chains (Fig. 4 bottom panels), as they loose direct contact during the opening motion. Residues in the RBD-S2 linker region in Chain A and C (residue 524-700) show a large gain in correlation in the initial stage of RBD opening (“slightly open”) despite being not directly in contact with the RBD in the closed form. On the pathway from close to open, relevant correlation changes are already found in the slightly-open state when comparing to the closed state (Fig. 4e), with significant changes for the important RBD-S2 linker region. However, more details appear (in other distant regions) as the transition approaches the partially-open and fully open states, cf. Fig 4f-g.

**Fig. 4.**
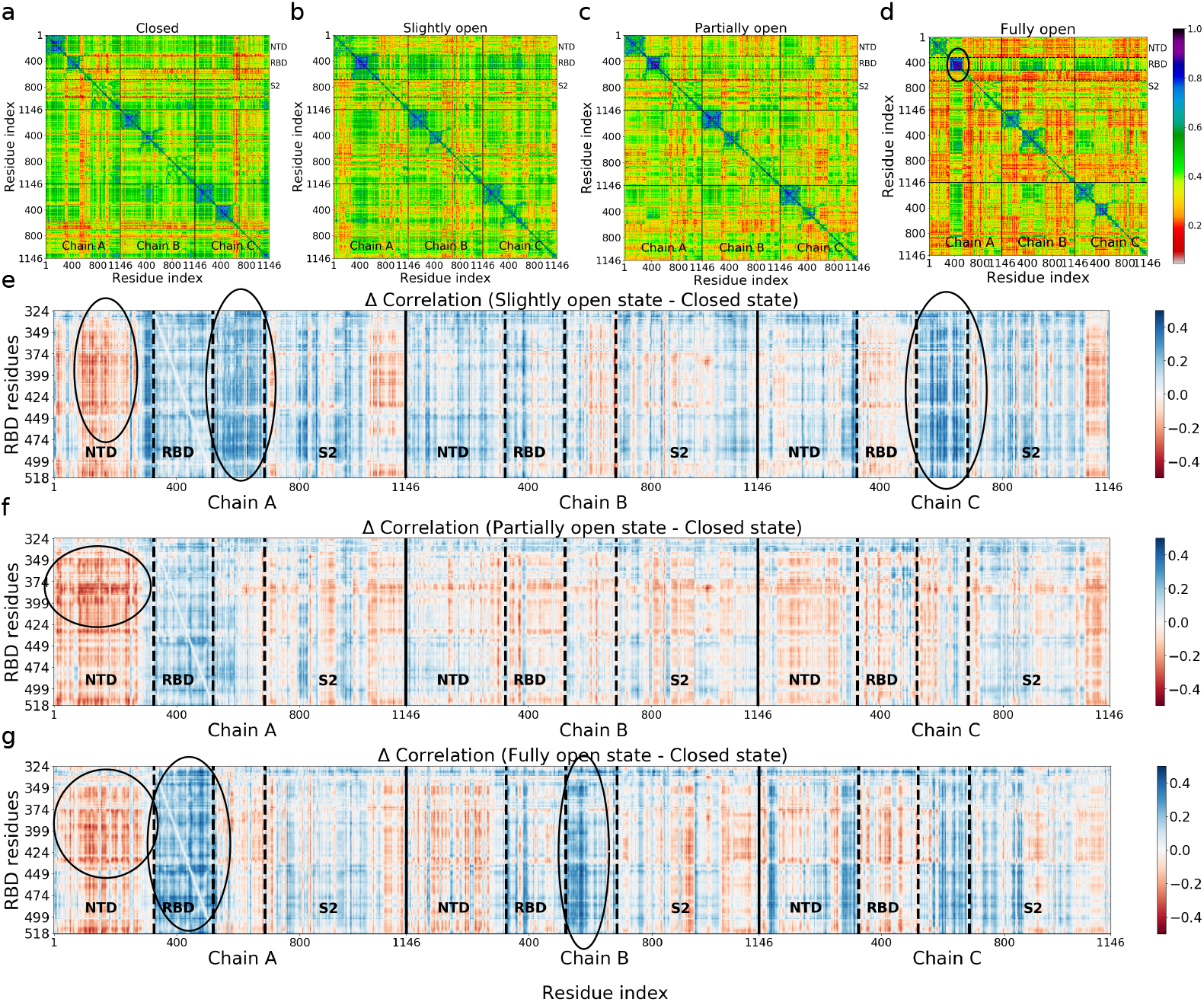
Upper panel: Cross correlation matrices for four states (a: closed, b: slightly open, c: partially open, d: fully open) computed using linear mutual information (LMI). Lower panel: Difference in correlation matrix elements for the (e) slightly, (f) partially and (g) fully open states with respect to the closed state. RBD regions form highly correlated blocks (d,g), indicating that these residues are largely decoupled from the rest of the protein. Still, signatures of long-distance correlated motion are detectable.

Overall these results are consistent with the dihedral angle correlations, described in the previous section: the residues in the loop region next to the RBD exhibit a change in the values of the backbone dihedral angles upon the down-to-up transition. The change in the correlation coefficient (ΔCorrelation) is also large for the RBDs and NTDs of chain B and C, which are in close proximity to chain A of the RBD in the closed state. Additionally, linker residues of chain B show significant gain in correlation upon the transitioning to the fully open state. Some residues that gain or loose correlation (blue or red coloration in Fig. 4 bottom panels) are situated at the opposite end (S2 region) of the spike, indicating the presence of long range correlated motion. This long-distance correlation can indeed be a cumulative effect of many small local fluctuations on the way towards the RBD, along structural patches connecting these sites, allowing “distant” residues to shed their impact on the structural transition in RBD. These pathways can be revealed, for example by the method of Ota and Agard (70), who used energy flow or vibrational energy relaxation to trace them.

For a more profound insight, we built a protein connectivity graph network-model. In it, the *C_α_* atoms of each amino acid are the nodes and the correlation between them are the edges connecting the nodes. The number of nodes in our system is N=3438, which makes it one of the largest systems studied previously with this method (62–67)(Comparable to the work by Saltalamacchia et al. on a splicosome complex involving 4804 *C_α_* atomd and 270 phosphorus atoms in the network (71)). We then calculated the betweenness centrality (BC), a graph theoretical measure that provides a way to quantify the amount of information that flows via the nodes and edges of a network. If a node *i* is working as a bridge between two other nodes along the shortest path joining them, then the BC of node *i* is given by

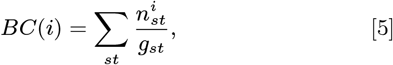

where *g_st_* is the total number of geodesics (shortest paths) joining nodes *s* and *t*, out of which 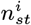 paths pass through node *i* (63). The change of BC in the dynamics of the spike protein has been recently observed using coarse-grained simulation methods(19). Despite the coarseness of the model, a handful of residues participating in the information-propagating pathway could be identified directly from the BC values. In the current work, we used the difference in BC as a metric to identify key residues which gain or loose relative importance along the allosteric information pathway. The difference in the normalized BC is measured by comparing the number for the partially open and fully open states with the closed conformation (i.e. *BC*^slightly open^ – *BC*^closed^ – *BC*^Partially open^ – *BC*^closed^ and *BC*^fully open^ – *BC*^closed^ for every residue in the spike protein) from our all atom trajectories with explicit solvation. Importantly, our model also includes the highly relevant glycan shield, which were shown to modulate the conformational dynamics of the RBD by favoring a *down* conformation and functioning as a gate for the conformational opening, beyond their general role in shielding (35, 38). However, glycans were not included in the network analysis. While in principle it is valid to consider their role, their motion occurs on time scales that are much faster than those of the protein backbone and would be averaged out of any correlation calculation.

For slightly open, partially open and fully open states, the residues with significant (e.g., >0.1) change in BC are mostly from the NTD region or RBD region of the B and C chain (Figs. 5 and 6). This suggests that the allosteric information flows through the nearby NTD’s and RBDs, and mutations in this region can break the allosteric network (63) and affect the functionality of the spike protein. SARS-CoV-2 neutralizing antibodies were indeed observed to bind these regions of the spike (22). The identified residues are in close proximity to the RBD of chain A, the one undergoing the *down* to *up* transition.

**Fig. 5.**
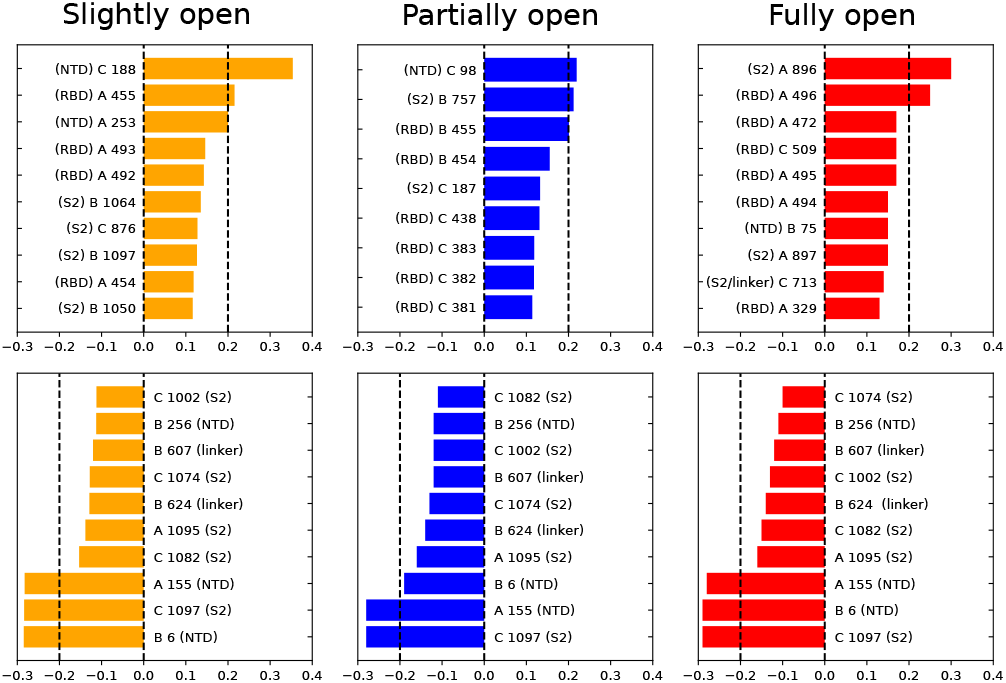
Residues with large changes in normalized betweenness centrality (BC) due to RBD opening. Upper panel: largest positive changes; lower panel: negative changes; chain index (A,B, or C) mentioned before the residue number. The location of the residue (NTD, RBD, linker or S2) is also mentioned next to the residue number.

**Fig. 6.**
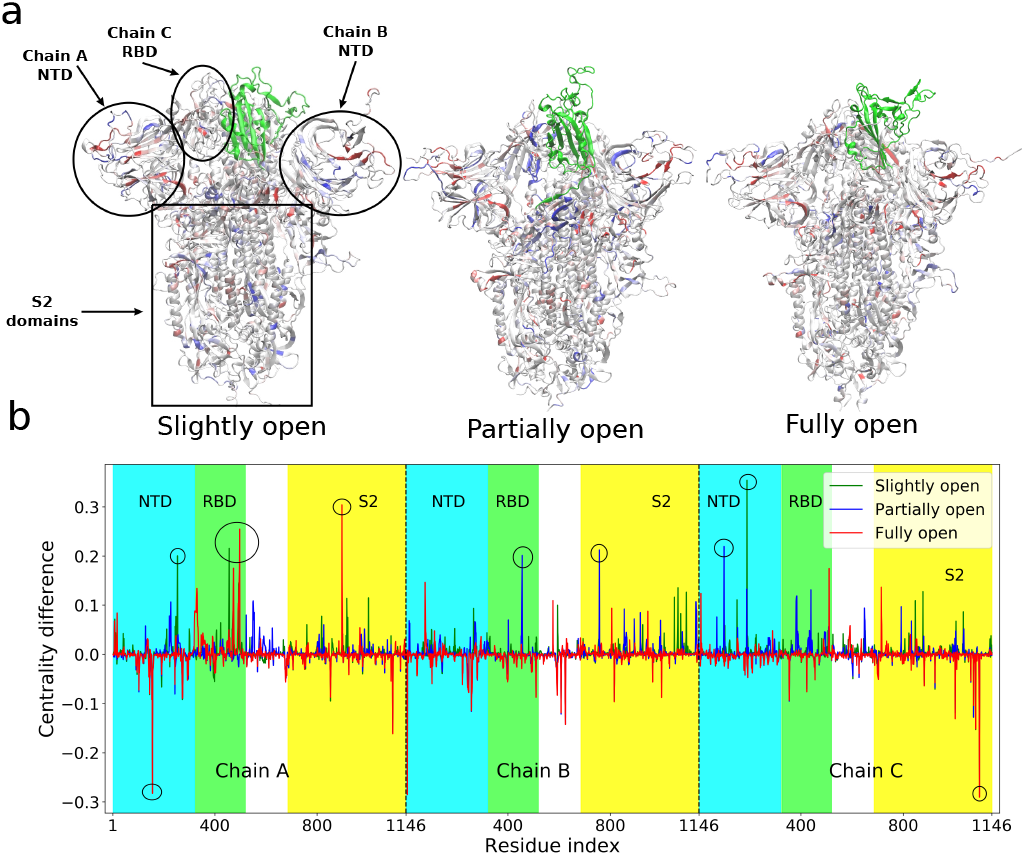
(a) Structures of the slightly open, partially open and the fully open states of the spike protein with residues colored according to the difference of betweenness centrality (BC) with respect to the closed conformation. Red indicates most negative and blue most positive values of BC difference. The RBD of chain A that undergoes opening motion is colored in green. The NTD and S2 domains are also indicated. (b) The values of the difference in centrality with respect to closed state plotted as a function of residue indices. Residues with BC difference > 0.2 marked with circle.

So the connectivity captured in the BC data is primarily due to short-range coupled fluctuations. Such couplings are broken when the RBD and NTD move apart, leading to the change in BC. For the same reason, the BC of the RBD of chain A increases in the fully open state as its internal vibrations become more independent of the rest of the protein.

In a culmination of the above, the most interesting aspect is the strikingly large change in BC of the residues which are *distant* from the RBD in 3D structure. Significant gain or loss of BC is observed in residues 607, 624, 713, 757, 896, and 1097. The first three residues are present in the linker region joining the RBD with the S2 domain, while the other three are in the S2 itself. The linker region has a strong impact on the dynamics of the RBD as we already established from the dihedral angle analysis. The allosteric network analysis reinforces this conclusion. Moreover, the large change of BC in the S2 domain indicates a complex long-range information flow connecting the RBD with the core residues of the protein. Electrostatic and van der Walls energy analysis, similar to that mentioned in the previous section, has been performed on the residues with changes in BC greater than 0.2. The interaction energy of the S2 domain residues such as I896 and G757 are significantly different for the closed and open state along some of the NTD residues like N188, V6, and S98. This has substantial implications for pharmaceutical design, as mutations within the NTD and the S2 domain can impact the receptor-binding propensity of the viral spike. These results also suggest that therapeutics targeted towards the S2 and towards the RBD-S2 linker can be effective in preventing COVID-19 infection, without complications stemming from the high rate of mutations in the RBD.

## Concluding Discussion

To tame the raging pandemic, we need to be able to control the fundamental dynamics of the spike protein. Its motion is key to the infection machinery of the SARS-CoV-2 virus.

By understanding the role of residues in its structure, we can anticipate the effect of new mutations and customize treatments ahead of time. To curb the negative impact of rapid mutations, we here focused on the allosteric effect of protein residues that are *away* from its rapidly-mutating epitope (the RBD) on the conformational change needed for infection.

We performed molecular dynamics simulations, with unsupervised machine learning (tICA) and graph theory-based analysis to identify the role of physically distant residues in the dynamics of the receptor binding domain in the SARS-CoV-2 spike protein. We correlated the protein backbone torsion angles with the slowest degrees of freedom encompassing the structural transition of the receptor binding domain. With this approach we were able to elucidate a small number of distant non-RBD residues which strongly influence the conformational change of the spike, change that in turn leads to binding to the ACE2 receptor and then to infection. Residues in the linker between the RBD and the S2 stem work as a hinge by driving the down-to-up RBD transition via backbone torsional changes. Out of the most correlated residues, D614 ranks close to the top. The D614G mutation is currently observed in SARS-CoV-2, and is becoming widespread among infected patients throughout the world. In vitro experiments also established that the single point mutation D614G is capable of altering the “down” to “up” conformational dynamics of the sike (55). The D614G mutant prefers the one RBD “up” state ~7 times more than the “3-down” configuration while they are equally likely in the wild type strain (54). The specific role of the D614G mutation was recently also established by coarse grained MD study(72). Our model predicted the D614 residue as a key player in RBD dynamics from physics-based atomistic simulations without any prior knowledge of the mutation profile of the spike. With glycine being more flexible in its backbone torsion compared to aspartate, this mutation, according to our hypothesis, will facilitate the attainment of the partially open state transiting from the closed structure. Another mutation, A570V, was observed within our predicted residues, but did not yet become as widespread as D614G, likely because it did not have substantial evolutionary advantage. However, a different mutation at residue 570 (A570D), has indeed appeared in the recently-emerged more-contagious B.1.1.7 strain of SARS-CoV-2 (41). The consistency with the mutation profile confirms that our dihedral-angle based analysis can not only find out distant residues impacting RBD dynamics, but can also predict residues where future mutations can increase infection capability.

A cross-correlation metric based on linear mutual information (LMI) was also employed to understand the long-distance coupled motion between RBD and non-RBD residues. The change in LMI correlation primarily takes place in the residues adjacent to the RBD, but we could also distinctly observe longdistance effects. Betweenness centrality (BC) of each residue of the spike was computed from a *C_α_*-based graph network model for all three conformational states. The residues showing largest changes in the BC are concentrated in NTD, RBD and also in the linker regions joining the RBD with the rest of the protein and also in the S2 domain. Dynamic allostery has been shown to impact the dynamics and consequently the binding strength of ACE2 receptor and antibodies to the mutated spike RBD (20). But significant change in centrality measures of the non-RBD domains in our study suggests that RBD dynamics is also impacted by long-distance allosteric effects within the spike protein itself; this emerges as a result of the collective internal fluctuations of the amino-acid residues. Experiments have not yet confirmed the role of S2 mutations for RBD opening dynamics and for its binding affinity to the ACE2 receptor. But whenever one observes a significant change in infectivity or virulence in the newly emerging strains of SARS CoV-2, it stems from a combined effect of multiple mutations in the S1 and S2 domain. As our computational approach has revealed that certain residues in the S2 domain can potentially modify the propensity of the conformational transition, the next logical step is to mutate those residues only and observe the effect, which hopefully will be addressed in future experimental studies. Moreover, the S2 domain is also involved in the “tectonic” conformational rearrangement required for the piercing of the host cell membrane (73), which we, however, do not study in the current work.

Although our theoretical predictions of relevant residues, connected to the changes in the RDB dynamics and effectively leading to higher virulence, can seem somewhat speculative, similar molecular dynamics studies on different proteins (74) could reveal single key residues in hinge domain driving major conformational changes, which subsequently have been confirmed by single-molecule experiments (75). We also point out that our simulation of the wild-type spike can, by fiat, only identify the residue to be mutated, but not the amino acid to which the residue will be mutated to. However, both experimental and computer simulation studies have already established that D614G, A570D and a few other mutations transform the dynamics of the RBD and favour an “up” state (54, 55, 59, 61, 72). In the current work, we highlighted a set of residues which show high correlation with RBD opening motion. We refrain from performing additional simulations by mutating those residues as our method cannot predict the end result of the mutation, and scanning over all possible mutations would be computationally expensive. But we do show that the dihedral angle preference, interaction energy and hydrogen bonding pattern of concerned residues change significantly due to a D614G mutation. Furthermore, the topic of whether the RBD opening motion increases infectivity can only be resolved *in vivo*, given the complexity of the viral entry process. A number of experiments and computational studies indicated that binding to ACE2 is feasible only when the RBD is “up” (27–31). This indicates that the down-to-up conformational change is indeed necessary for binding to ACE2. While we did not thoroughly study the dynamics of actual mutant spike proteins, the role of specific point mutations in the dynamics of RBD of the spike protein has been explored using MD simulation in the literature. (60, 72)

From the point of view of immediate therapeutic interventions, this study opens up the possibility of designing inhibitors that bind to the regions outside the RBD, thereby preventing infection by freezing RBD dynamics via steric restrictions on specific distant residues. Such treatments are less likely to be affected by the evolutionary adaptations in RBD sequence that the virus performs frequently to evade the immune response. In a starker context, future mutations in these key residues can potentially change the infection rate and virulence, giving rise to new strains and significantly altering the course of the pandemic. Our study and future work in this direction can make the scientific community better prepared for such scenarios and can help in efficient prevention of future outbreaks.

## Data availability

The molecular dynamics trajectories are available from http://doi.org/10.5281/zenodo.5052691. The codes and the residue correlation data used in this study are available from https://github.com/dhimanray/COVID-19-correlation-work.git. All further details about the methods and the data are available within the article and the SI Appendix.

## Materials and Methods

The details of molecular dynamics simulations, tICA analysis, and mutual information based network analysis are provided in the SI Appendix. A brief outline is included below.

### System preparation and simulation details

Glycosylated and solvated structures of closed (PDB ID: 6VXX (32)) and partially open (PDB ID: 6VSB (34)) spike head trimers were obtained from the CHARMM-GUI COVID-19 archive (76). The glycans included in the simulation are those from Table 1 of Ref (76). All simulations were performed using the CHARMM36m force field (77). For the purpose of the current work we considered residues 324-518 as the RBD. After minimization and short equilibration, steered molecular dynamics (SMD) simulation were performed to induce the opening of the closed state and closing of the partially open state of the RBD. Reaction coordinates (RC) were chosen to represent the distance of the RBD from its closed state position. Multiple structures were chosen from the two SMD trajectories and two independent umbrella sampling (US) simulations were performed for PDB ID: 6VXX and 6VSB. The reaction coordinate was restrained by a harmonic potential with force constant of 1 kcal/mol/Å^2^ for 35 windows and 26 windows for the two sets respectively. The colvars module (78) was used for SMD and US calculations. Free energy profiles were computed using weighted histogram analysis method (WHAM) (79).

### Unbiased simulations

Umbrella sampling trajectory frames were sampled from the regions near the open, closed and the partially open intermediate state judging the free energy value. Unbiased simulations were performed starting from these frames, resulting in 39 trajectories, each 40 ns long. Three of these trajectories were identified as stable conformations corresponding to the closed, partially open and fully open structure. These three trajectories were extended to 80 ns. A cumulative ~ 1.7 *μ*s unbiased simulation data were generated and used in subsequent analysis. Additional 40 ns simulation was performed for the D614G mutant spike for each of the closed and the partially open state. The structures of the mutated species were generated using UCSF Chimera package (80).

### Time-lagged independent component analysis and mutual information

time-lagged independent component analysis (tICA) and principal component analysis (PCA) were performed using pyEMMA package (81) on the entire unbiased trajectory data. The feature space for PCA and tICA consisted of pairwise distances between specific residues in and around RBD of chain A and the NTD and core domains.

The linear mutual information (LMI) based correlation was computed for the closed, slightly open, partially open and fully open state trajectories. A graph theory based network model was constructed with the *C_α_* atom of each residue as node. The edge length between nodes were computed from the cross correlation values using previously described procedure (62). Betweenness centrality of each residue was computed for each of the three trajectories and compared. All the LMI and network analysis were performed using bio3D package (82).

### Sequence alignment

Iterative sequence alignment of the 67 strains of SARS-CoV-2 spike protein sequences from the RCSB PDB database was performed using the MAFFT-DASH program (83) using the G-INS-i algorithm. The sequence of PDB ID: 6VXX was used as the template. The alignment was analyzed with the ConSurf server (84) to derive conservation scores for each residue position in the alignment.

## ACKNOWLEDGMENTS

This work was supported by the National Science Foundation (NSF) via grant MCB 2028443. DR acknowledges support by the Molecular Science Software Institute (MolSSI) seed COVID-19 fellowship funded by NSF via grant number OAC-1547580. The authors thank Trevor Gokey for a stimulating discussion. The work has benefited from the computational resources of the UC Irvine High Performance Computing (HPC) cluster.

