## Supplementary material for "Distant Residues Modulate Conformational Opening in SARS-CoV-2 Spike Protein": SI Appendix

#### This PDF file includes:

Supplementary text
Figs. S1 to S17 (not allowed for Brief Reports)
Table S1 (not allowed for Brief Reports)
SI References

### Supporting Information Text

### Additional results

**Free energy profile of RBD opening.** We performed extensive umbrella sampling simulation to obtain the underlying free energy landscape of the *down* to *up* transition of the receptor binding domain. Roy et al. observed that this transition in SARS-CoV-2 spike is a complex multidimensional process with unique intermediates different from SARS-CoV-1 and MERS (1). They characterized an unusual Proline-proline interaction (P230-P521) between the RBD and the N-terminal domain which stabilizes the intermediate partially open aka "NTD-assisted up" state. In consistency with their finding, our enhanced sampling simulations could not distinguish between three different states based on single geometric reaction coordinate. Following the work by Folding@Home group (2), our first set of simulations starting from the RBD down structure (PDB ID: 6VXX) used the distance of the center of mass of the RBD from the same in the closed state, as the RC. We could obtain only one deep free energy minima which corresponds to the closed state (Fig S1a). The position of the minima is in agreement with the most probable structure predicted in Ref (2). The small 3 Å separation of the free energy minimum from the cryo-EM structure is the representative of the most stable state for the given force field. The free energy profile is virtually identical to the results of Ref (1) when RBD-NTD interactions were artificially removed. It indicates the all atom enhanced sampling simulation starting from RBD down state is unable to reach the partially open configuration within reasonable amount of computational time. To sample the conformations around the NTD-assisted up state of the RBD, we performed another set of enhanced sampling simulation starting from the experimental structure of the partially open intermediate (PDB ID: 6VSB). The underlying free energy profile qualitatively resembles the coarse grained MD results of Ref (1) but with fewer approximations. Unlike the previous work (1) our free energy profile, along a inter-residue distance based RC, indicates a lower free energy of the partially open state compared to states which are closer to the RBD-down configuration (Fig. S1b). This is not entirely unexpected as the closed-like state obtained from the second set of simulations does not relax to the true closed state, indicated by insufficient increase of the P230-P521 distance. Also the non-equilibrium work profile shows a very high value (20 kcal/mol) for the open conformation, which is much higher than the equilibrium free energy of that configuration (Fig S1a). This is not a problem as obtaining the true underlying free energy landscape is not the goal of the present work. We used the umbrella sampling trajectories to seed multiple unbiased simulations at and near the closed, partially open, and fully open states.

**Results for D614G mutant.** We performed additional simulations on the D614G mutant spike protein to verify our predictions about the dihedral angle preference and interaction energy. The $\phi$ angle of residue 614 and the $\psi$ angle of residue 613 show strong correlation with the RBD opening motion in the wild type strain and occupy different regions of the torsion angle configurational space in the closed and partially open state. But in D614G mutant, the distribution of both these angles overlap during the course of the simulation, irrespective of being in closed or open state (Fig S14). This reinforces our hypothesis that the D614 residue, which is predicted to play a key role in the dynamical transition in the spike protein from our model, no longer show different dihedral angle conformation preference between RBD down and RBD up state, when mutated to glycine. There is a significant contribution arising from the electrostatic energy as well. In the partially open state of the wild type spike, there is an additional peak in the electrostatic energy distribution of D614 residue with the rest of the system. The peak is more than 100 kcal/mol higher in energy (less negative) compared to the most probable electrostatic energy of the closed state. This difference is reduced to 20-30 kcal/mol upon the D614G mutation (Fig S15). This is not surprising because the absence of the charged side chain prevents the formation of stabilizing interactions like hydrogen bonds and salt bridges in the mutant. It is pronounced by the fact that, unlike the wild type system, the number of hydrogen bonds forming between the residue 613-614 remains unchanged between the closed and the partially open states in case of the D614 mutant (Fig. S16).

**Sequence alignment and analysis.** We performed sequence alignment of the 67 sequences of the SARS-CoV2 spike protein from the Protein Data Bank. Analysis of the alignment using the ConSurf server (3) indicates that most residues of the spike protein are highly conserved. In agreement with previous studies, (4, 5) the variable residues are primarily concentrated in the receptor binding domain (Fig. S17). This result motivated the current work focusing on the role of non-RBD residues in the infection process in search of long term targets for therapeutic design. Glycosylations (6) are typically found in the conserved regions of the protein, (except for N1134) to shield these regions from the host immune system.

At the end of our study we found that all of the strongly correlated residues predicted from our dihedral angle analysis are highly conserved (Gln613, Pro600, Gly601, Asn137, Ser112, Cys136, Phe833, Ile834, Ile569, His1083, Asp1084), except for Asp614 and Ala570, mutations in both of which have led to new, highly contagious, strains which we discussed extensively in the main text. Most of the non-RBD residues involved in the allosteric pathway of RBD opening are also highly conserved. The high level of conservation reinforces their important role in the functioning of the spike as well as the possibility of targeting them using drugs and antibodies to obtain broad spectrum inhibition of the spike.

### Methods

**A. System preparation and equilibration.** The cryo-EM structures of the SARS-CoV-2 spike (S) protein in closed (PDB ID: 6VXX (7)) and partially open state (PDB ID: 6VSB (8)) were used in this work. Fully glycosylated and structurally complete (missing residues rectified) S protein head-only models (residue 1-1146) were obtained from the CHARMM-GUI COVID archive (9). The two protein structures were obtained as pre-solvated in cubic water box of edge length 201 Å and 202 Å respectively. The ion concentration was maintained at 150 mM by including appropriate number of $K^+$ and $Cl^-$ ions in the system. The total number of atoms in the closed and partially open S-protein systems were ~762,000 and ~773,000 respectively. All proteins and glycans were modelled using CHARMM36m (10) force field and a TIP3P model was used for the water.

Molecular dynamics simulations were performed using NAMD 2.14b2 package with CUDA acceleration (11). Each structure was first minimized using conjugate gradient algorithm for 10000 steps which was followed by a short equilibration in NPT ensemble for 250 ps with 2 fs time step. The temperature were controlled using a velocity re-scaling thermostat and a Langevin barostat was used to control the temperature and pressure at 310.15 K and 1 atm respectively. The systems were further equilibrated at the same temperature and pressure for 10 ns with hydrogen mass re-partitioning (HMR) (12) where the mass of each hydrogen atom was made 3 a.m.u. and the mass of the connecting heavy atoms were adjusted to keep total mass unchanged. This allowed for the use of 4 fs time-step to integrate the equations of motion. The thermostat was switched to a Langevin thermostat with 1/ps damping coefficient. The equilibrated structures obtained at the end were subjected to further study. The simulation protocol for the rest of the work was identical to the second round of equilibration (except for the application of harmonic biases in umbrella sampling which is mentioned in the respective section). Up to this point, all simulation input files were obtained from CHARMM GUI COVID-19 archive and were used without modification. They are freely available in the CHARMM-GUI web-server (http://www.charmm-gui.org/?doc=archive&lib=covid19) and can be accessed for further simulation details.

**B. Steered molecular dynamics and umbrella sampling.** The open structure and multiple intermediate structures for the spike protein were generated by steered molecular dynamics. The RBD deviation coordinate (2) (the distance of the center of mass of RBD residues in a given frame of simulation from the RBD center of mass in the crystal structure) was used as a reaction coordinate (RC) to perform SMD on the closed structure (6VXX). For the other system (6VSB), the RC was chosen to be the distance between Asp428 residue of chain A (the one whose RBD is going from “down" to “up" conformation) and the Lys986 residue of chain C. These residues are in proximity in the closed form and are far apart in the partially open and open conformation. For the SMD trajectory starting from the closed (PDB ID: 6VXX) system, the value of the RC was varied from 0 to 35 Å at constant velocity over 8 ns of simulation. A moving harmonic restraint was employed on the RC with a force constant of 1 kcal mol$^{-1}$ Å$^{-2}$. The system was kept at the final state with a static restrain for additional 2 ns. All protein $C_\alpha$ atoms, except for the ones pertaining to residues 300 to 600 in chain A, were harmonically restrained with 0.5 kcal mol$^{-1}$ Å$^{-2}$ force constant. This prevents any spurious conformational change of the rest of the spike protein during the application of strong steering force on the RBD of Chain A. For the system generated from the partially open form (PDB ID: 6VSB), SMD simulation was performed with identical condition except the RC was varied from 35 Å to 10 Å over the first 8 ns.

Structures, which were ~1 Å apart in RC space, were sampled from the SMD trajectories to perform umbrella sampling. For the first set (originating from PDB ID: 6VXX) 35 structures were sampled between RC value 1Å to 35Å. For each structure, a 17 ns simulation was performed in NPT ensemble. The restraint on the RBD-deviation coordinate was gradually increased to 1 kcal mol$^{-1}$ Å$^{-2}$ over the first 100 ps, and was held constant for the rest of the simulation. The trajectory data of the last 11 ns was used for the analysis. For the second set (originating from PDB ID: 6VSB) 26 structures were sampled for the RC (Asp428(A)-Lys986(C) distance) values between 10Å to 35Å. Each trajectory was subjected to 15 ns of simulation with harmonic restraint on the RC, and last 10 ns of each trajectory was used for the analysis. Rest of the protocol is identical to the first set. Weighted Histogram Analysis Method (WHAM) (13) was used to calculate the 1D PMF from both set of trajectories. Error bars were computed from the Monte Carlo estimator implemented in WHAM program (14).

**C. Unbiased simulation and time-lagged independent component analysis.** Multiple unbiased trajectories, each of length ~40 ns were initiated from the last frames of specific umbrella sampling simulations. Particular care was practiced in the choice of the starting structures. The US windows corresponding to 1Å to 10Å can better represent the vicinity of the closed state, while RBD-deviation of 22Å to 30Å corresponds to fully open structures, as realized from manual inspection of the trajectories. The partially open state is supposed to be the intermediate between closed and fully open states. So we chose umbrella windows 16Å to 35Å from the second set (starting from PDB ID: 6VSB) of US simulations to represent the conformational space in and around the partially open state. As all of the trajectories, intended to sample the fully open conformations, showed gradual RBD closing motion, we performed an additional simulation starting from the 33Å window of the first set. In total 39 unbiased simulations were performed. The trajectories corresponding the umbrella window 3Å and 33Å of the first set and the 32Å from the second set were identified as the closed, fully open and partially open state judging from the free energy profile. These unbiased trajectories were extended to ~80 ns. A cumulative ~ 1.7 $\mu$s of unbiased simulation data was generated.

The unbiased trajectories were subjected to principal component analysis (PCA) (15) and time-lagged independent component analysis (tICA) (16, 17). First, all trajectories were projected onto a feature space consisting of contact distances between pairs of residues from the RBD and the rest of the spike protein. Particularly, we focused on the residue pairs which are close to each other in the closed state and far in the open state. We identified residues within the region 250-550 of chain A, we identified residues whose $\alpha$-carbon is within 8 Å of the $\alpha$-carbon of any other residue in the rest of the spike. The $C_\alpha - C_\alpha$ distances of such pairs (a total of 173 distances) were used as the feature space for PCA and tICA analysis.

As we found that the tICA analysis can better distinguish between all three states, only tICA results were used for subsequent analysis. The projection of the trajectories in the space of first two principal components has been depicted in Fig. S2. We computed the sines and cosines of the backbone dihedral angles ($\phi$ and $\psi$) of all residues in the spike and their Pearson correlation coefficients with the two slowest degrees of obtained from tICA analysis. The residues were ranked based on the correlation score ($CS$) metric described in the main text.

**D. Mutual information and cross correlation.** The four trajectories corresponding to the closed, slightly open, partially open and fully open states, were used to calculate mutual information and inter-residue cross correlation. This protocol is traditionally used for calculating allosteric connections in Apo and Holo protein ligand systems. We followed this protocol except we used four different conformational states of the spike protein.

The theoretical details of this method are described in Ref (18–20). In information theory the mutual information metric $(I_{ij})$ between pairs of atoms can be computed as

$$I_{ij} = \int\int p(\mathbf{x}_i, \mathbf{x}_j) \ln\left(\frac{p(\mathbf{x}_i, \mathbf{x}_j)}{p(\mathbf{x}_i)p(\mathbf{x}_j)}\right) d\mathbf{x}_i d\mathbf{x}_j, \qquad [1]$$

where the $\mathbf{x}_i, \mathbf{x}_j$ are the vectors containing the Cartesian coordinates of the atoms, whereas $p(\mathbf{x}_i)$ and $p(\mathbf{x}_j)$ are their marginal distributions and $p(\mathbf{x}_i, \mathbf{x}_j)$ is the observed joint distribution (20). A Pearson like correlation coefficient $C_{ij}$ can be obtained from $I_{ij}$ using the following expression:

$$C_{ij} = (1 - e^{-(2/d)I_{ij}})^{-1/2} \qquad [2]$$

We used linear mutual information in our analysis. It is given by

$$I_{ij} = H_i + H_j - H_{ij} \qquad [3]$$

where $H$ is a Shannon type entropy function:

$$H_i = -\int p(\mathbf{x}_i) \ln p(\mathbf{x}_i) d\mathbf{x}_i$$
$$H_{ij} = -\int\int p(\mathbf{x}_i, \mathbf{x}_j) \ln p(\mathbf{x}_i, \mathbf{x}_j) d\mathbf{x}_i d\mathbf{x}_j \qquad [4]$$

$\mathbf{x}_i, \mathbf{x}_j$ are the vectors containing the Cartesian coordinates of the $C_\alpha$ atoms.

The difference between the cross correlation heat map is computed to highlight the difference of long distance interactions between the RBD and the rest of the protein in the slightly open, partially open and fully open states in comparison to the closed state. Despite the approximate nature of the artificially generated "fully open" state we did include this trajectory for mutual information calculation. The reason behind that is when we are building a model to predict the effect of individual residue in a conformational transition we aim to include as much trajectory data of different configurations as possible. As it is computationally expensive to compute the cross correlation map for such a large system using all 39 trajectories, we only chose four conformationally distinct states that can possibly be involved in the transition.

We further constructed a protein graph connectivity network from the mutual information correlation matrix for each of the three states. We calculated and compared the between centrality (BC) for each residue. The BC provides a way to quantify the amount of information flow between nodes or edges of a network. As the $\alpha$-carbon of each residue is considered to be a node in our network construction, the value of BC provides the functional relevance of a residue in the total information flow through the protein. If a node $i$ is working as a bridge between two other nodes along the shortest path joining them, the BC of node $i$ is given by

$$BC(i) = \sum_{st} \frac{n_{st}^i}{g_{st}}, \qquad [5]$$

where $g_{st}$ is the total number of geodesics (shortest paths) joining nodes $s$ and $t$, out of which $n_{st}^i$ paths pass through node $i$ (21, 22). We measured the change in BC for each residue by quantifying the difference their values between two open states and the closed state.

**Analysis of hydrogen bond and non-bonded energies.** The number of hydrogen bonds for each residue group with from table 1 (main text) were computed using VMD H-bond toolkit. The default criteria of a donor acceptor distance of 3.0 Å and an angle cutoff of $20^o$ was used. The number of hydrogen bonds in all frames of the closed and partially open state trajectory was obtained and the mean and 95% confidence interval was computed. The electrostatic and van der Walls energies of residues with high CS score (from Table 1 main text) and for residues with BC difference greater than 0.2 were computed using the NAMD Energy plug-in of VMD software. Both hydrogen bonding analysis and the non-bonded energy analysis were performed on the closed state and partially open state trajectories.

**E. Simulation of D614G mutant spike protein.** To justify our predictions about the possible future mutants we performed additional simulation for the D614G mutant spike protein. The first frame of the longer equilibrium simulations for closed state and the partially open state were used to construct the initial structures for the corresponding states for the D614G mutant. The mutations were performed in all three chains using UCSF Chimera package (23). Glycans, identical to the wild type system, were added to the mutant spike using CHARMM GUI. The systems were solvated and ionized to have 0.15M KCl concentration. Conjugate gradient minimization was performed for 10000 steps followed by 1 ns NPT equilibration with 2 fs timestep and harmonic restrain on all $C_\alpha$ atoms to preserve the structure. Production runs were performed for 40 ns for each system starting from the end point of the equilibrated trajectory. All other parameters were identical to the simulations of the wild type systems.

 

**F. Sequence alignment and analysis.** We performed sequence alignment for 67 sequences of SARS-CoV2 Spike Protein from Protein Data Bank. The software for constructing the alignment is MAFFT-DASH (24). MAFFT utilises FFT algorithm to accelerate iterative sequence alignment without sacrificing too much accuracy. The web-based DASH (Database of Aligned Structural Homologs) provides structural alignments at the domain and chain levels for all proteins in the Protein Data Bank (PDB), resulting in improvement for alignment involving multiple sequences with weak similarity. The G-INS-i algorithm was used with maximum iteration cycles of 1000. This algorithm assumes that entire region can be aligned and tries to align them globally using the Needleman-Wunsch algorithm; that is, a set of sequences of one domain must be extracted by truncating flanking sequences. The result of sequence alignment is shown in figure 1, with sequence of PDB ID: 6VXX as the template. The alignment was analyzed by ConSurf server (3) to derive conservation score for each residue position in the alignment. The input for ConSurf analysis includes the sequence alignment by MAFFT above, atom coordinate of 6VXX and the sequence of 6VXX as the template. The residues will be sorted by a scale of 9 levels of conservation, starting from very high variability to very high conservation. The distribution of conserved and variable amino acids on the structure of S protein can be visualized by mapping the conserved and variable residues are also mapped onto the structure of 6VXX taken from PDB (Fig. S17). The mapping tool was also provided on ConSurf server. The map shows that conserved residues are predominant throughout the structure of S protein whereas most variable residues cluster around the tip of the receptor binding domain.

**Table S1. A list of simulations performed in the current study**

| Simulation type | Starting point | Number of trajectories | Length of each trajectory (ns) |
|---|---|---|---|
| SMD | Equilibrated closed structure (PDB: 6VXX) | 1 | 10 |
| SMD | Equilibrated partially open structure (PDB: 6VSB) | 1 | 10 |
| Umbrella Sampling (US) | Points sampled from SMD trajectory (PDB: 6VXX) | 35 | 17 |
| Umbrella Sampling (US) | Points sampled from SMD trajectory (PDB: 6VSB) | 26 | 15 |
| Unbiased MD | End points of selected US trajectories (PDB: 6VXX)<br>RC=1Å,2Å,4Å-10Å,22Å-30Å, with 1Å interval | 18 | $\sim$ 40 |
| Unbiased MD | End points of selected US trajectories (PDB: 6VSB)<br>RC=16Å-19Å,21Å-31Å,33Å-35Å with 1Å interval | 18 | $\sim$ 40 |
| Unbiased MD (Closed) | End point of US trajectory (PDB: 6VXX)<br>for RC=3Å | 1 | $\sim$ 80 |
| Unbiased MD (Partially open) | End point of US trajectory (PDB: 6VSB)<br>for RC=32Å | 1 | $\sim$ 80 |
| Unbiased MD (Fully open) | End point of US trajectory (PDB: 6VXX)<br>for RC=33Å | 1 | $\sim$ 80 |
| Unbiased MD (Closed)<br>D614G mutant | End point of US trajectory (PDB: 6VXX)<br>for RC=3Å and mutated | 1 | $\sim$ 40 |
| Unbiased MD (Partially open)<br>D614G mutant | End point of US trajectory (PDB: 6VSB)<br>for RC=32Å and mutated | 1 | $\sim$ 40 |

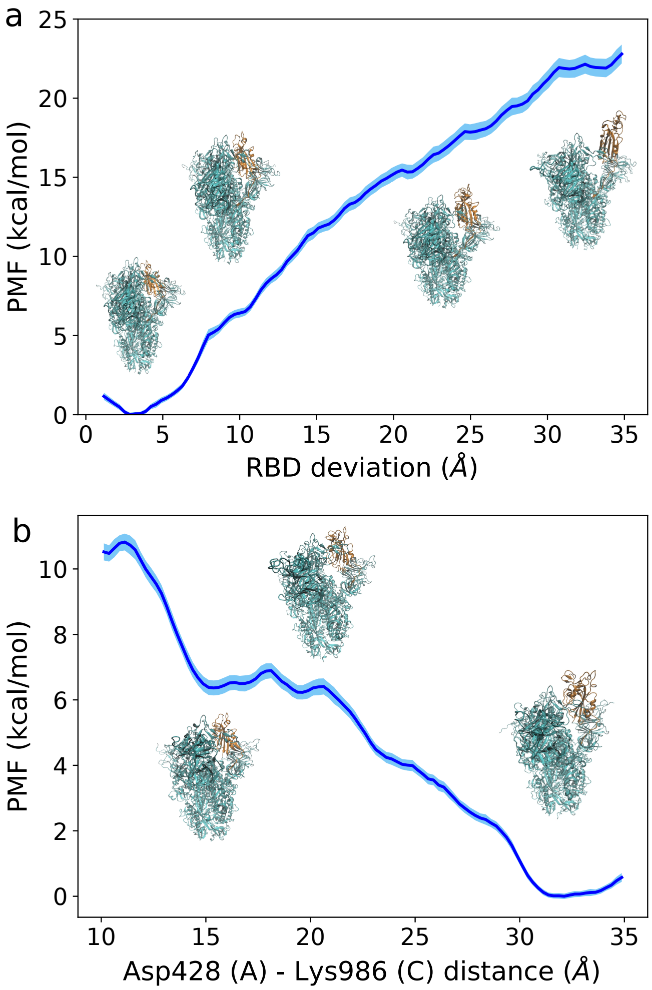


**Fig. S1.** Free energy profile along the RBD opening coordinate for (a) Set 1 (starting from PDB ID: 6VXX) and (b) Set 2 (starting from PDB ID: 6VSB). For further details see SI Methods section and SI text. Representative structures are shown along on the PMF. The transiting RBD has been colored in orange whereas the rest of the spike is colored light blue.

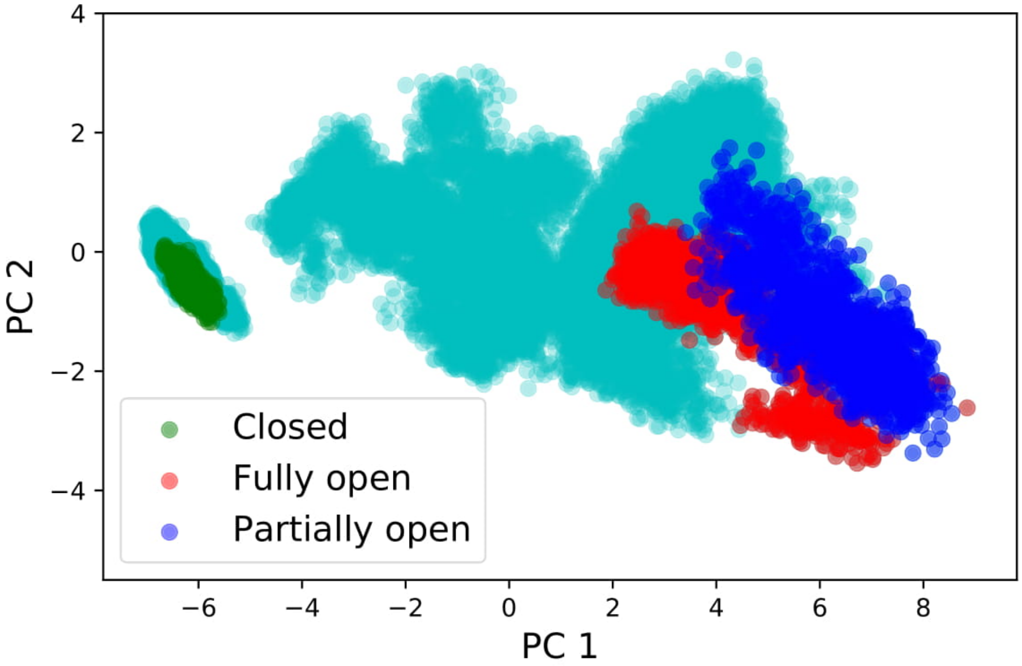


**Fig. S2.** The projection of all unbiased trajectories along the first two principal components obtained from PCA.

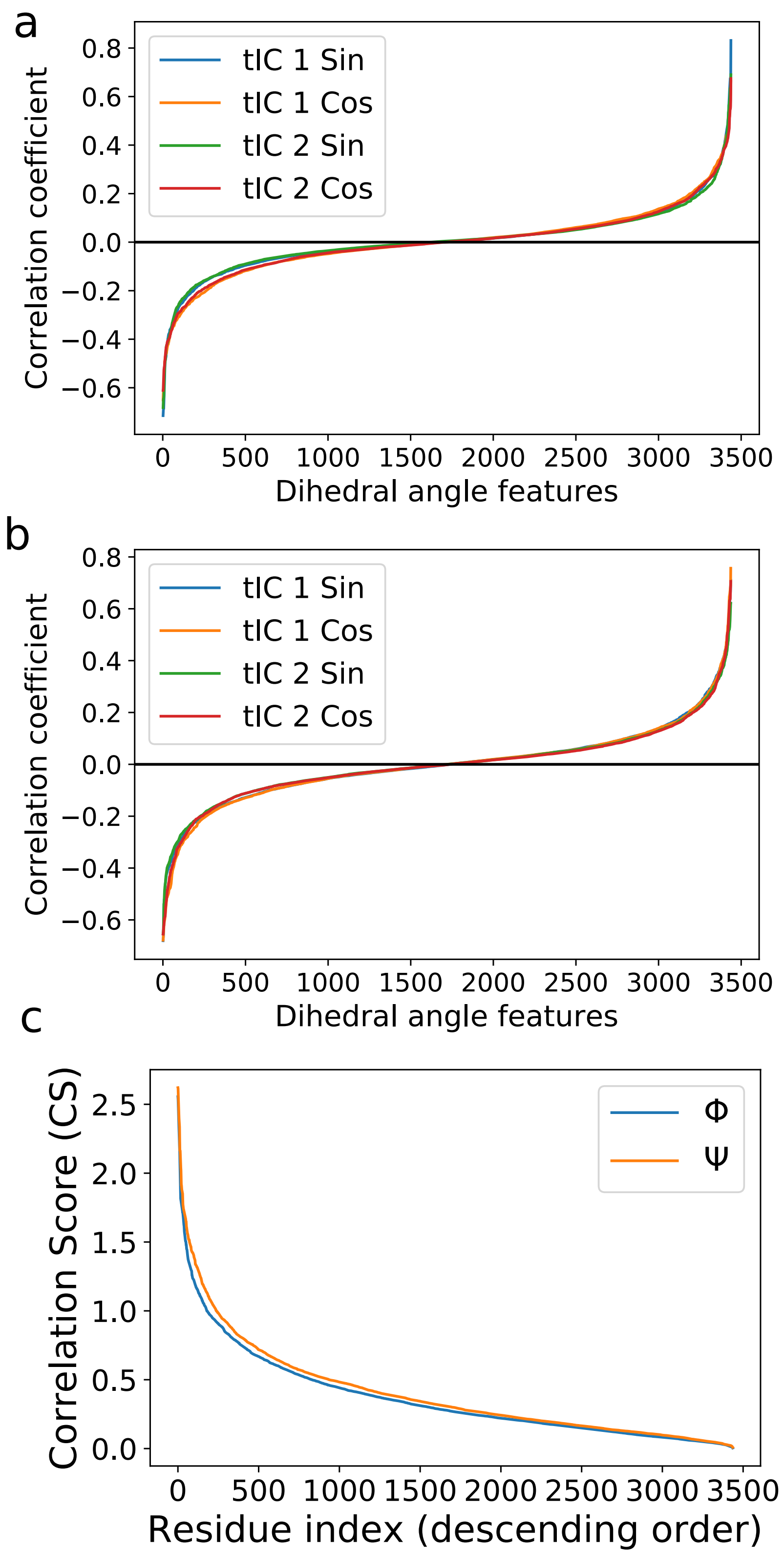


**Fig. S3.** (a) Correlation coefficients of the $\Phi$ angles of all residues arranged in ascending order shows that only a handful of residues have high positive or negative correlation with either two tICs. (b) same a (a) but for $\Psi$ angles. (c) CS scores of all residues in descending order indicating that small number of residues have high correlation score.

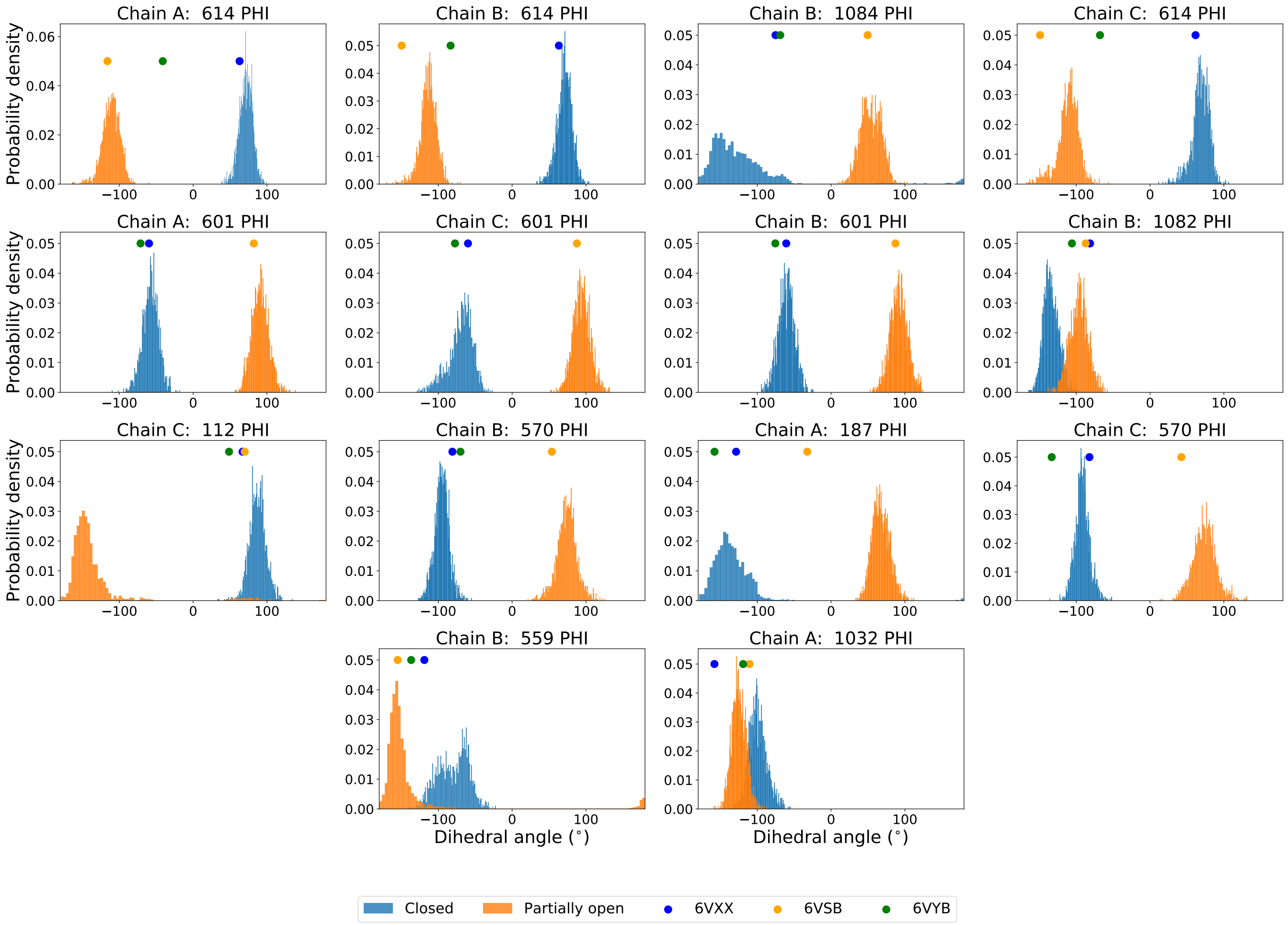


**Fig. S4.** Distribution of the dihedral angles, the **sine** functions of which are most **negatively** correlated with **tIC 2**. The distribution is plotted for closed, fully open and partially open state trajectories. The value of torsion angles in PDB structures a shown in dots.

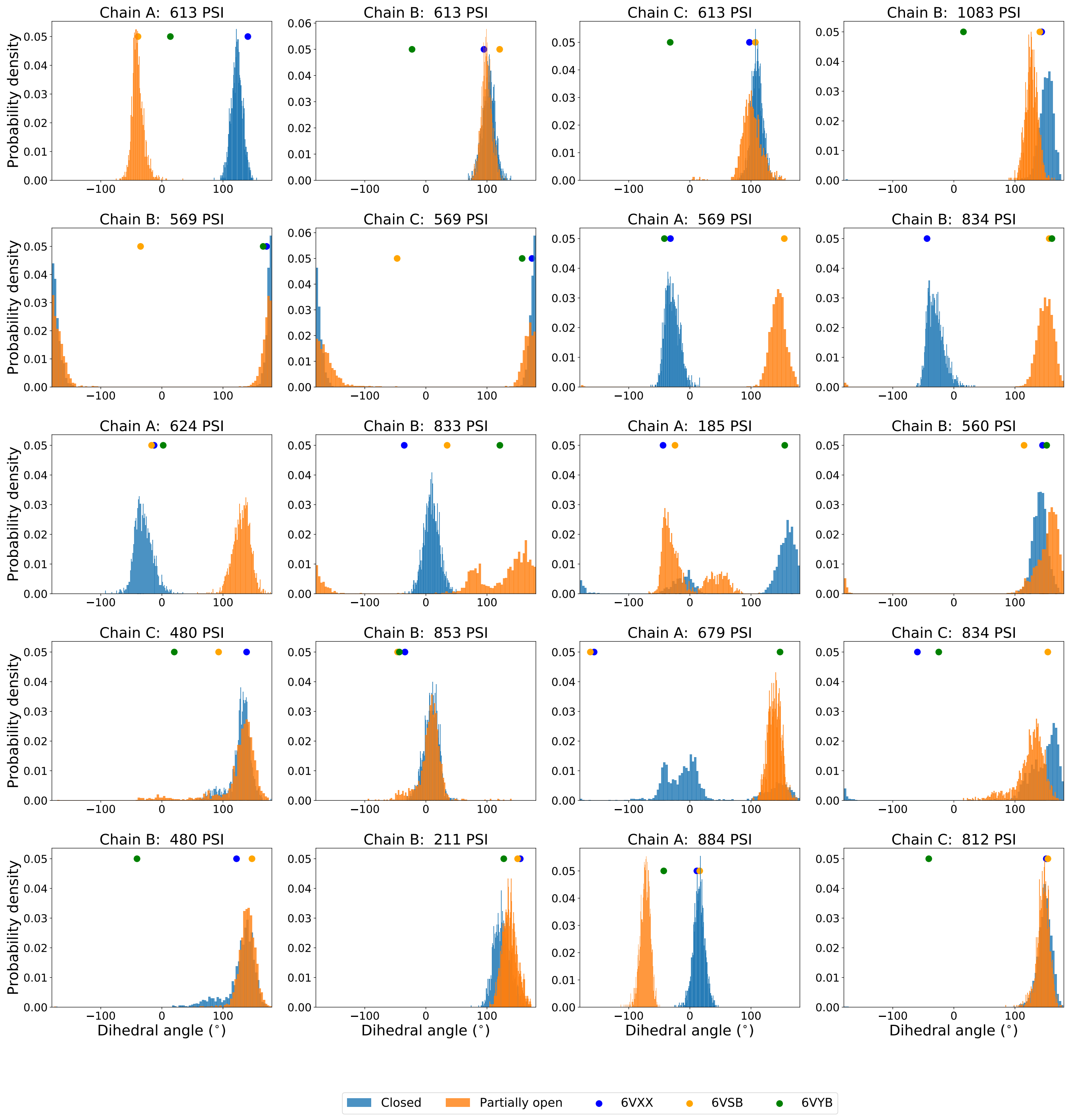


**Fig. S5.** Same as Fig. S4 except for **positive** correlation

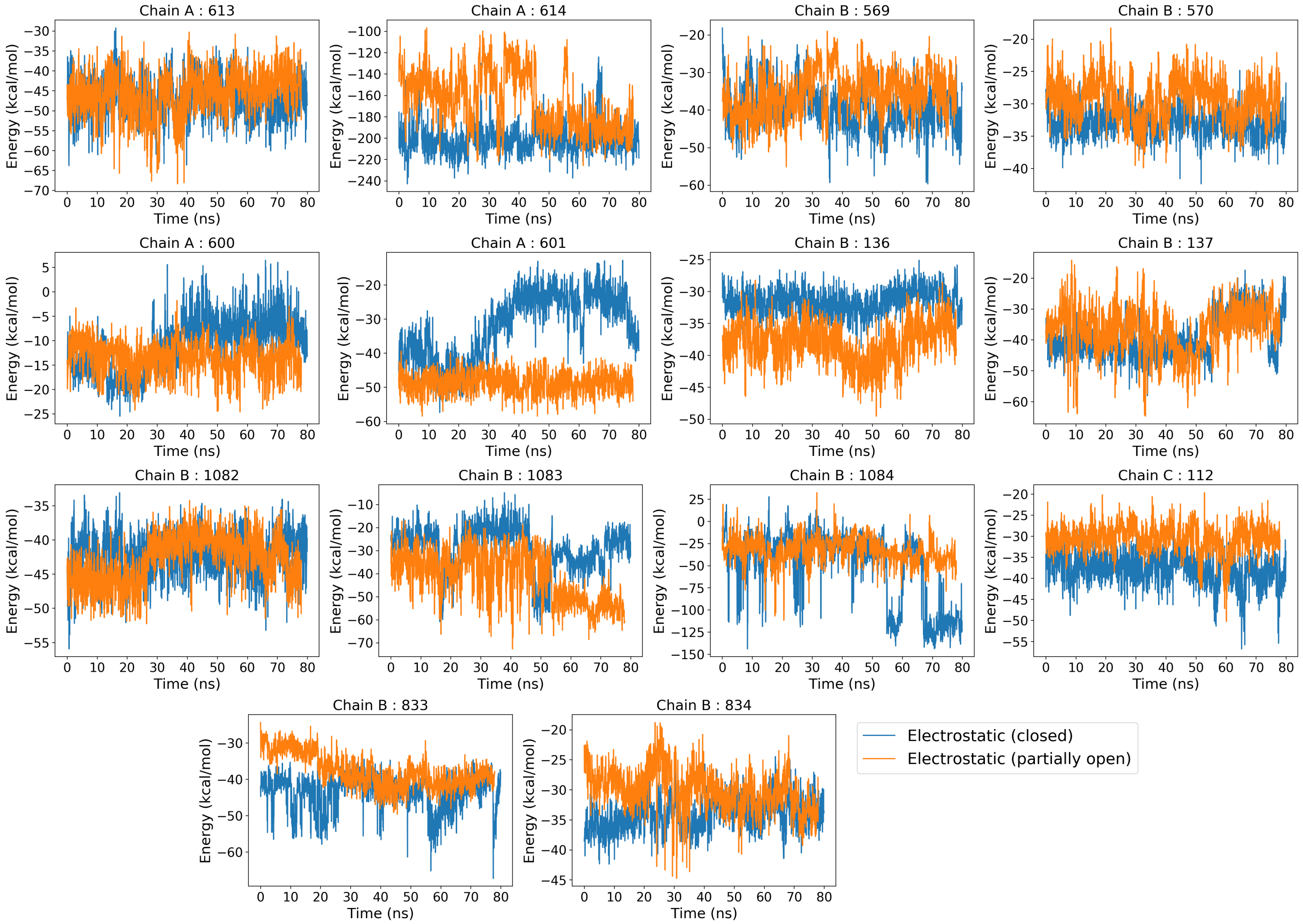


**Fig. S6.** Fluctuation of electrostatic energy as a function of time for some of the residues with highest correlation score from tICA analysis. The results are plotted for the closed and the partially open state.

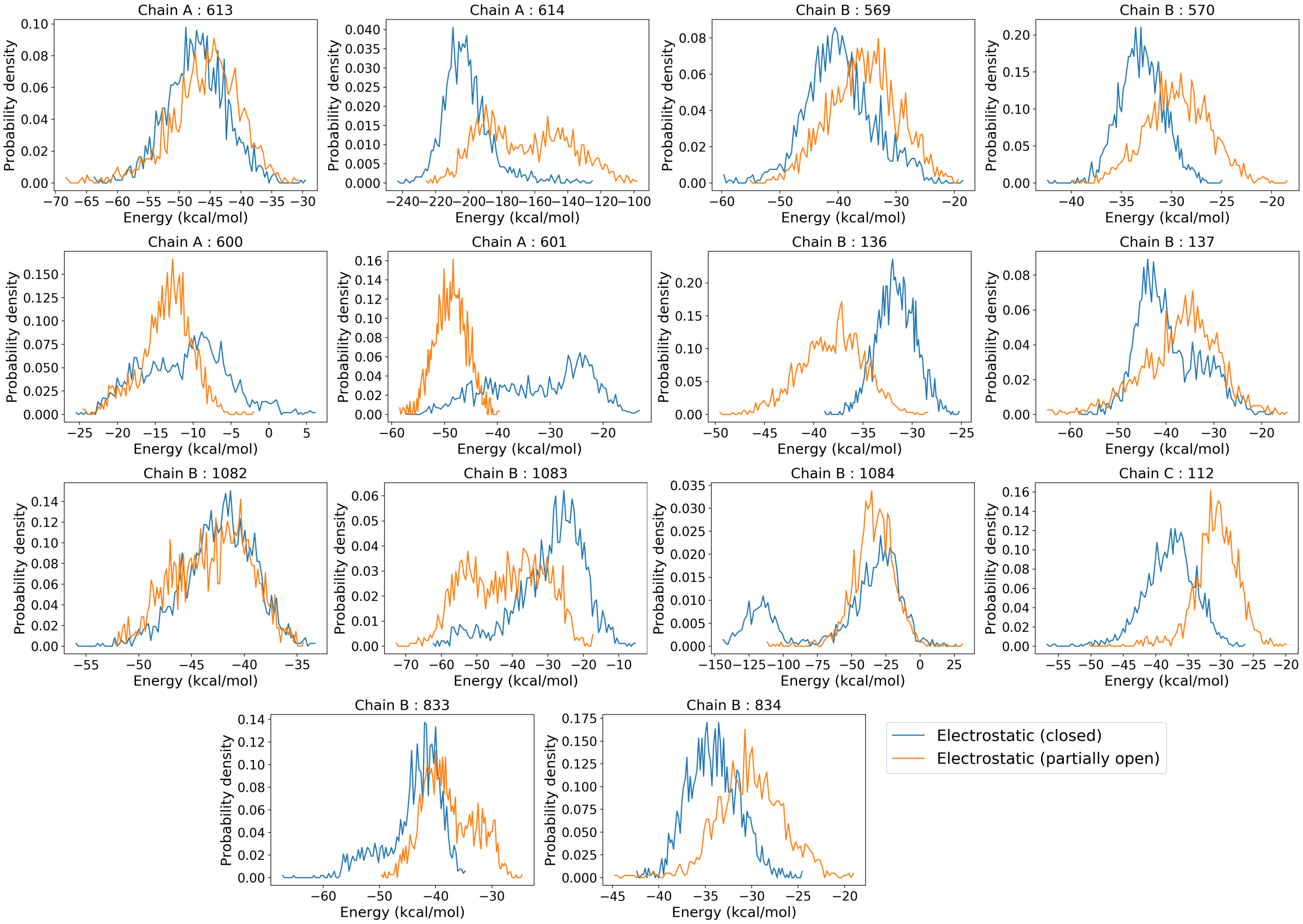


**Fig. S7.** Probability distribution of the electrostatic energies of residues from Fig. S6

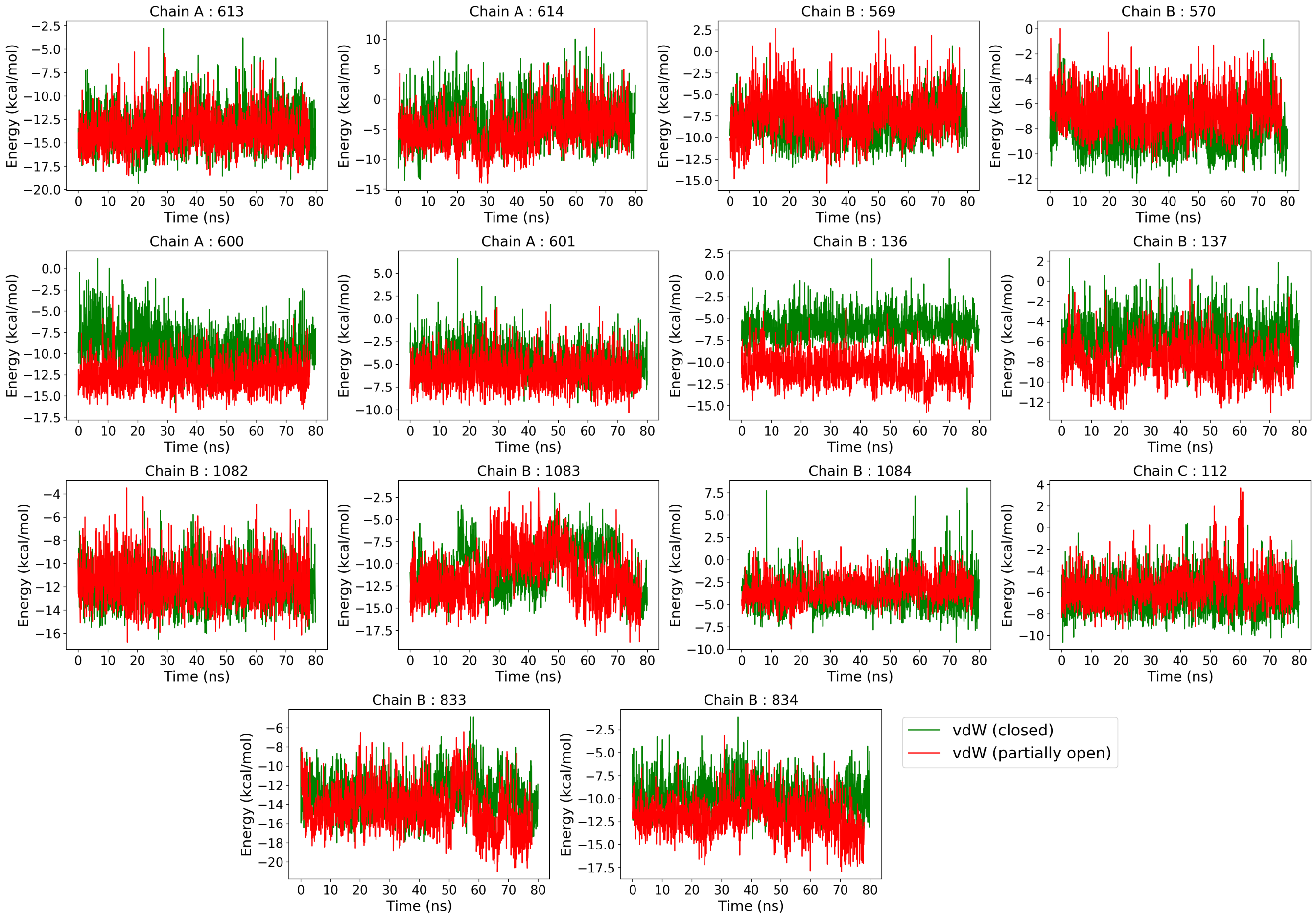


**Fig. S8.** Same as Fig. S6 but for van der Waals energies

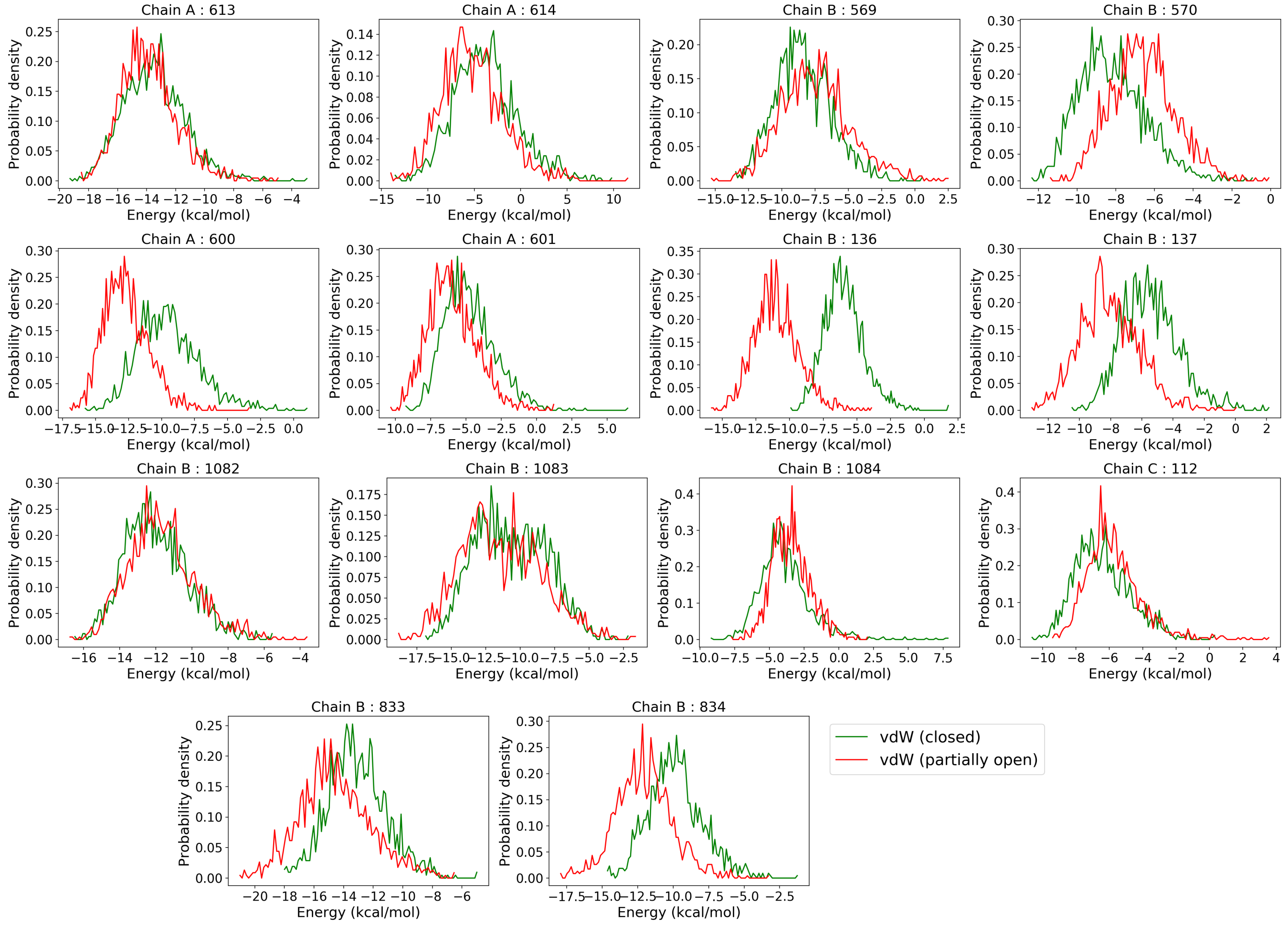


**Fig. S9.** Same as Fig. S7 but for van der Waals energies

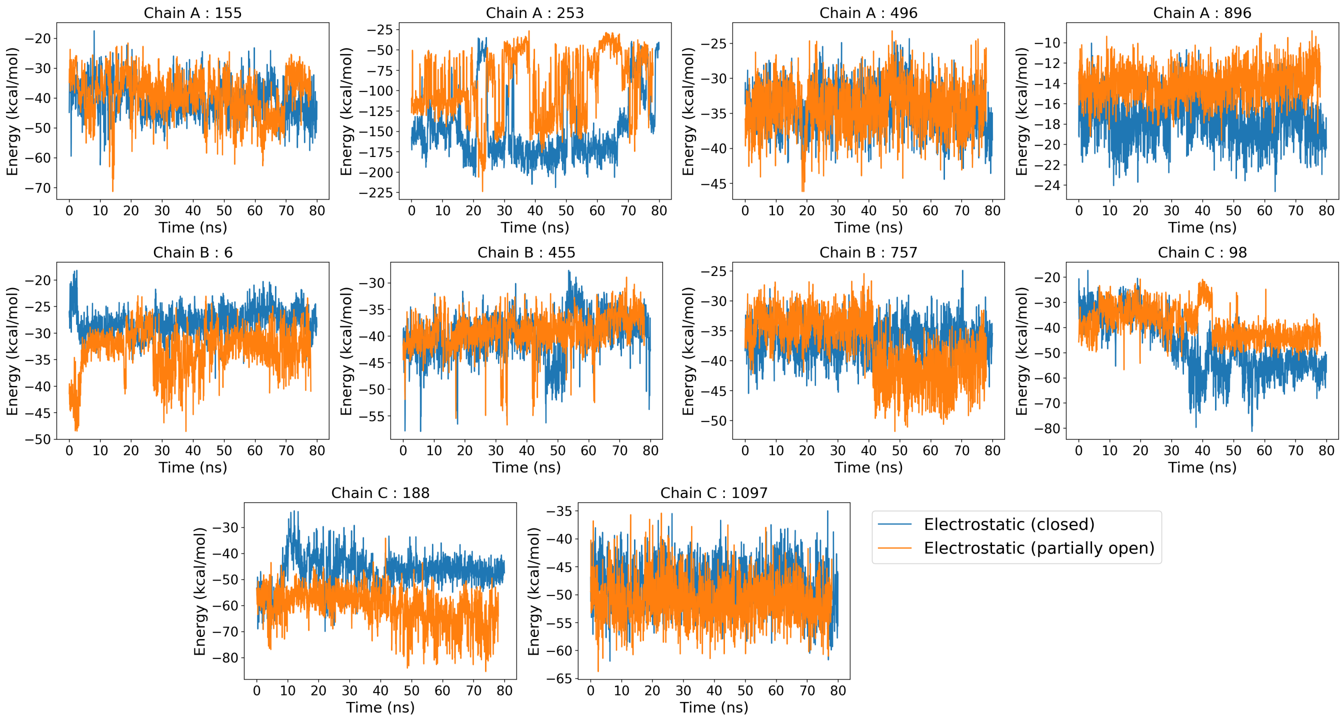


**Fig. S10.** Fluctuation of electrostatic energy as a function of time, for the residues with highest change in betweenness centrality (BC) upon RBD opening. The results are plotted for the closed and the partially open state.

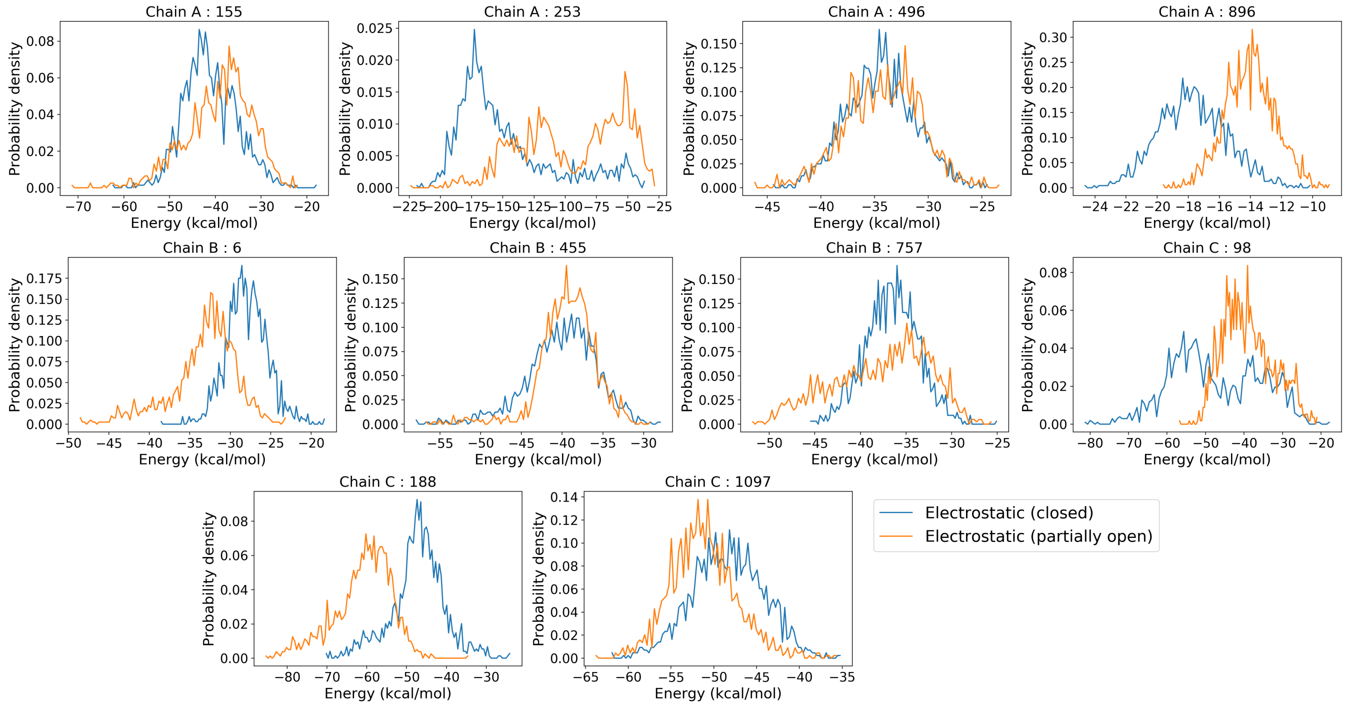


**Fig. S11.** Probability distribution of the electrostatic energies of residues from Fig. S10

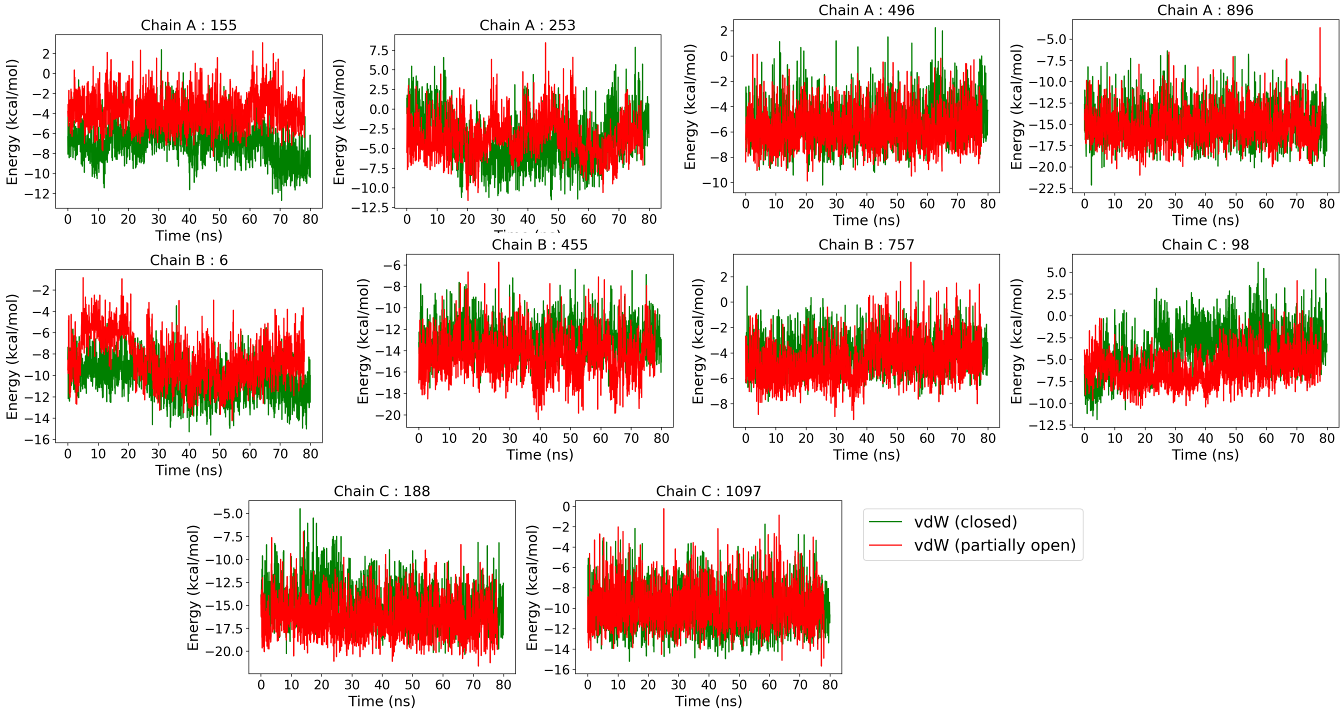


**Fig. S12.** Same as Fig. S10 but for van der Waals energies

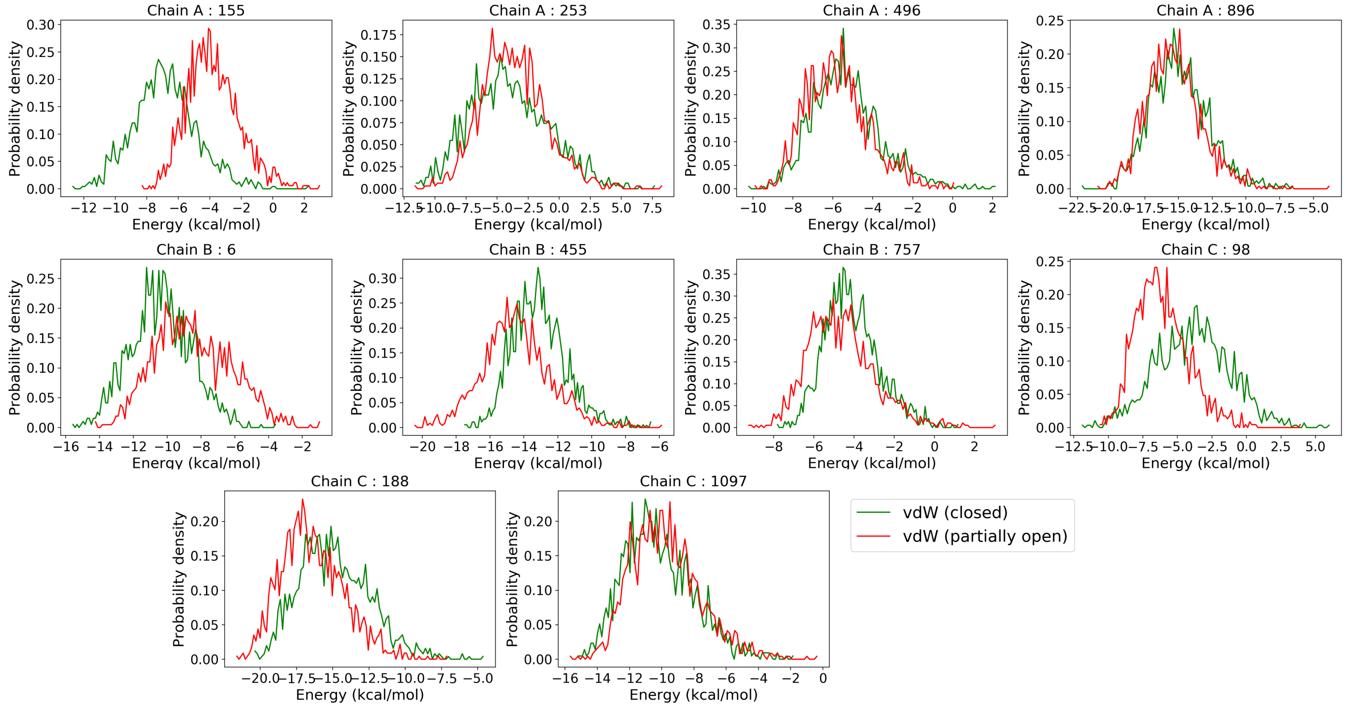


**Fig. S13.** Same as Fig. S11 but for van der Waals energies

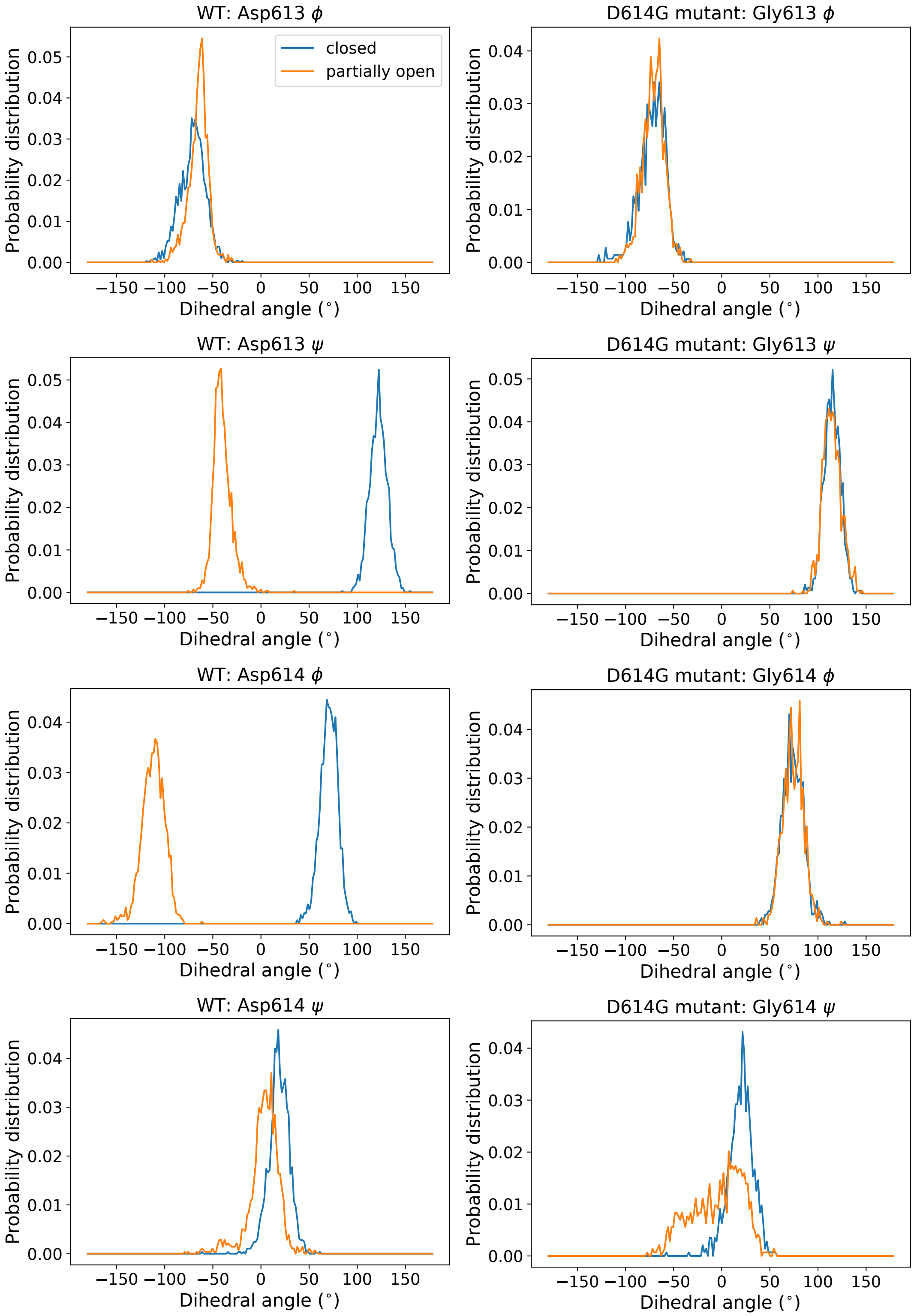


**Fig. S14.** Distribution of the backbone torsion angles of residue 613 and 614 for the wild type spike protein and the D614G mutant spike protein. The D613($\psi$) and D614($\phi$) angles occupy distinct regions in closed and partially open state in the WT system. But the A613($\psi$) and A614($\phi$) torsion angles show identical values for both states in the mutant.

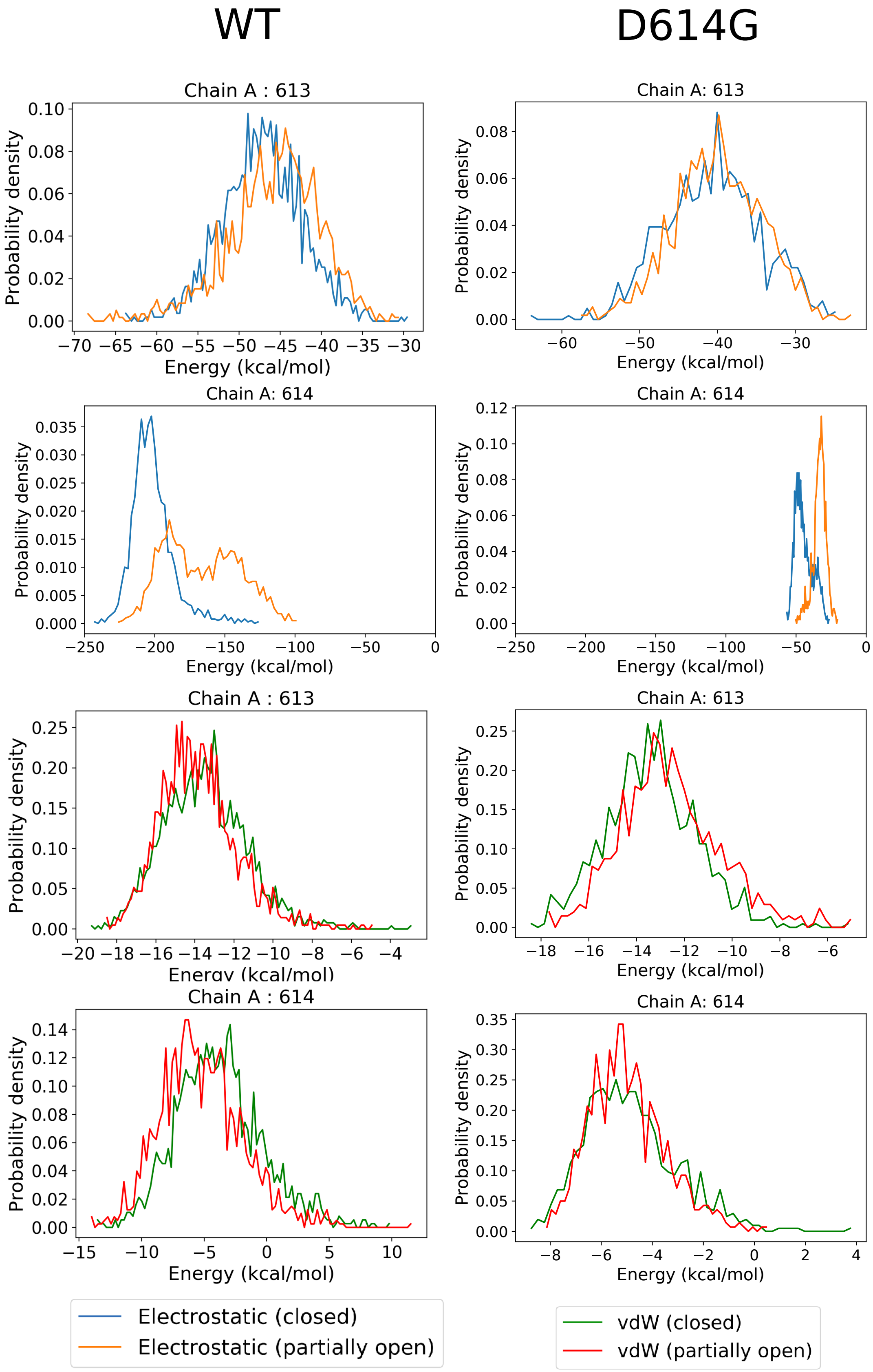


**Fig. S15.** Distribution of the electrostatic and vdW interaction energies of the residue 613 and 614 for the wild type spike protein and the D614G mutant spike protein.

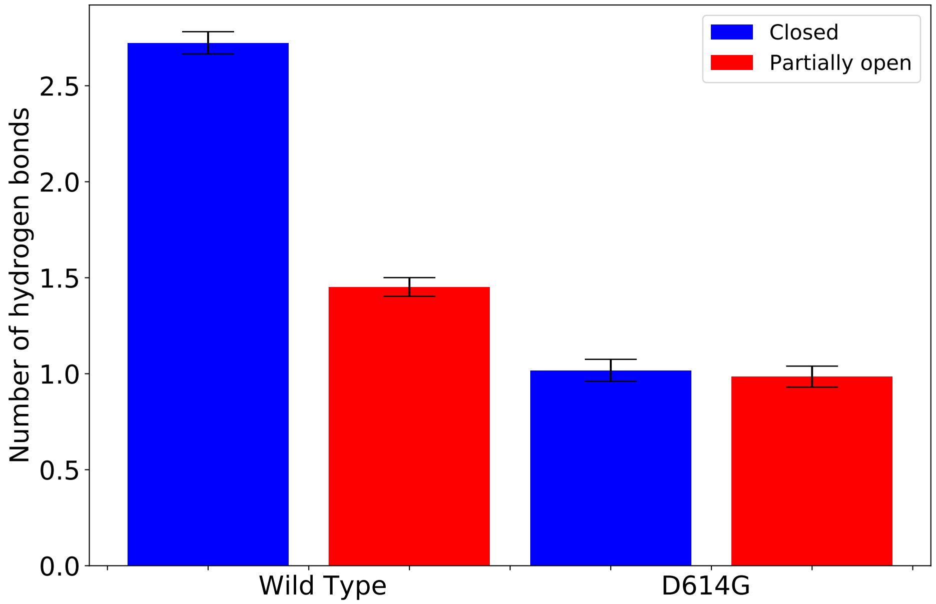


**Fig. S16.** The number of hydrogen bonds formed by the residue 613 and 614 combined, for the wild type spike protein and the D614G mutant spike protein.

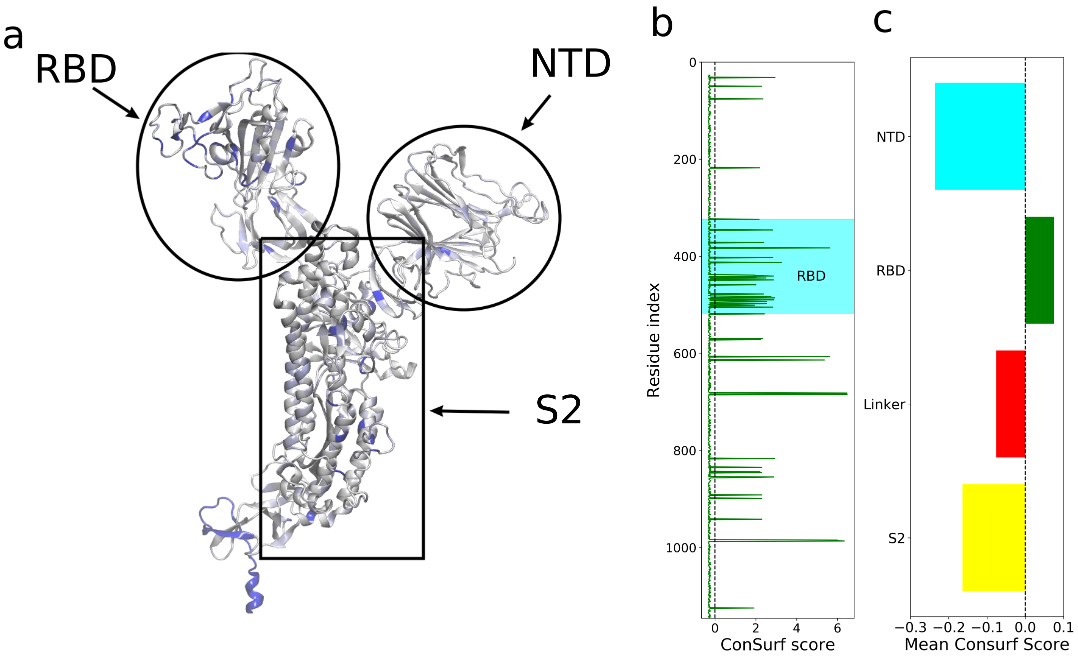


**Fig. S17.** (a) The structure of the spike monomer with residues colored according to their degrees of variability over different strains. The blue indicates most variable regions and white color shows conserved regions. (b) The ConSurf score for all residues. The region corresponding to the RBD is denoted in cyan. The lower (more negative) the ConSurf score, the more conserved are the residues, and vice versa. (c) Mean Consurf Score for each of the four domains indicating RBD is the most variable region, as it is the only domain with mean positive ConSurf score
